# Graph-Based EEG Symmetry Features from the Temporal Lobe as Markers of Antidepressant Treatment Response

**DOI:** 10.64898/2026.01.26.701781

**Authors:** Akbar Davoodi, Ioannis Vlachos, Martin Bareš, Martin Brunovský, Milan Paluš

## Abstract

Major Depressive Disorder (MDD) is a mental disorder that affects millions globally and has highly individualized responses to antidepressant treatment. Identifying objective early markers that can distinguish responders from non-responders remains a critical challenge in personalized psychiatry. In this study, we investigate the role of brain symmetry in EEG-derived functional connectivity as a potential early marker of treatment response. Using resting-state EEG recordings from 176 patients diagnosed with MDD at baseline (Visit 1) and after one week of treatment (Visit 2), we construct graph-based representations of functional connectivity and define region-wise and electrode-wise temporal-lobe symmetry features. We focus on symmetry *change scores* (Visit 2 − Visit 1) across typical EEG frequency bands in both weighted and binary forms.

Statistical analyses reveal a consistent group-level pattern in the *α* and *β* bands: responders show negative symmetry change scores, whereas non-responders show relative stability or weakly positive change scores. The contrast is strongest in the *β*_1_ band and is more pronounced for binary symmetry metrics, while the *α* band shows the same direction with slightly weaker significance. Physiologically, these opposite trajectories are *consistent with* a treatment-related shift toward more lateralized temporal-lobe network organization in responders and a more bilateral coupling pattern in nonresponders; however, these neurophysiological interpretations remain hypothesis-generating given the resting-state design and the absence of direct behavioral or task-based validation.

To quantify whether temporal symmetry change scores contain reproducible discriminative signal under strict validation, we benchmark the proposed features in a supervised classification pipeline using repeated nested cross-validation. The best-performing configuration achieved a median out-of-sample AUC of 0.690 with a 95% bootstrap confidence interval of [0.611, 0.711], indicating modest but reproducible separability.

Overall, temporal-lobe symmetry change features provide interpretable candidate markers of early antidepressant-related neurophysiological change that may complement existing EEG predictors in future multi-marker models, but they are not yet sufficient for standalone clinical decision support and require external validation in independent cohorts.

## Introduction

Major Depressive Disorder (MDD) is a widespread and serious mental health condition affecting an estimated 300 million people worldwide (1–3). The World Health Organization has recognized MDD as a major public health concern, projecting it to become the leading cause of disease burden by 2030 (4). This stark reality highlights the urgent need for effective treatment strategies and tools that can help predict who will benefit from specific interventions, helping individuals receive the right care at the right time.

One of the most challenging aspects of treating MDD is that people respond very differently to the same treatments. Some patients find significant relief with standard therapies, while others continue to struggle despite various treatment attempts. This variability underscores the importance of early and accurate predictors of treatment response. If clinicians could predict early on who is likely to benefit from a particular antidepressant, they could make timely adjustments, sparing patients the distress of prolonged ineffective treatments. Electroencephalography (EEG) is a commonly used tool in the study of the brain and has emerged as a promising tool in the field of MDD research (5). Non-invasive and relatively affordable, EEG captures millisecond-level brain signals, giving insights into the brain’s complex activity, interconnections and dynamics. This makes EEG particularly useful for studying mental health conditions like depression, where the brain’s functional networks can reveal clues about emotional and cognitive states.

In recent years, EEG has helped MDD researchers investigate brain connectivity, how different regions of the brain communicate with each other, especially in relation to predicting treatment outcomes for depression (6, 7). Previous studies have shown that connectivity strengths from certain brain areas can indicate MDD severity and even predict treatment responses (8). However, a less explored aspect is the role of brain symmetry and asymmetry. Simply put, brain symmetry refers to the balance or similarity of activity across corresponding regions in both hemispheres, while asymmetry refers to specialized processing in one hemisphere more than the other. Differences in symmetry, particularly in regions like the frontal and temporal lobes, could potentially explain why some patients respond better to treatment than others.

Brain asymmetry has long been linked to emotional regulation, with specific patterns reflecting either resilience or vulnerability to stress and depression. Although much of the earlier work concentrated on frontal asymmetry (9), recent neuroimaging and electrophysiological studies have increasingly highlighted the role of the temporal lobe in emotional processing, memory, and social cognition (10–12). For example, Toga and Thompson (11) mapped functional asymmetries that suggest the temporal regions contribute critically to balanced emotional states, while Hugdahl (10) and Rogers and Vallortigara (12) have argued that lateralized processing in these regions may underlie differences in mood and stress reactivity. These findings collectively support the notion that temporal lobe activity, and by extension its symmetry, could provide important insights into the neural mechanisms of Major Depressive Disorder (MDD) and its response to treatment.

While our focus on temporal lobe symmetry represents a novel approach, we acknowledge the substantial body of work examining other regional markers. Resting frontal alpha asymmetry (FAA) has long been investigated in MDD (5, 13), however, recent meta-analyses and large-scale replication attempts indicate limited and heterogeneous diagnostic value, with the largest datasets failing to confirm a robust FAA-MDD association (14–16). In contrast, converging evidence across PET/fMRI and source-localized EEG suggests that higher pretreatment activity, often theta, within the rostral/subgenual anterior cingulate cortex is a reproducible, treatment-nonspecific prognostic marker of symptom improvement across multiple interventions (17–19). Here we extend this literature by focusing on temporal-lobe network symmetry derived from imaginary coherence graphs as an interpretable, connectivity-level marker that is complementary to both frontal FAA accounts and cingulate-centric predictors.

From a clinical neuroscience perspective, hemispheric specialization in temporal regions underpins adaptive emotional regulation. The right temporal cortex predominates in processing negative emotions and autonomic arousal, whereas the left temporal lobe supports approach-related, positive affective states (20, 21). Antidepressant treatment is thought to rebalance these dynamics, restoring normative lateralization in networks critical for emotional memory and social-emotional decoding (10–12). Accordingly, we hypothesize that responders will show a treatment-related increase in temporal asymmetry, reflecting renewed functional specialization, whereas non-responders may manifest persistent or compensatory bilateral engagement. This is the main hypothesis of this work that we test and later interpret in the Discussion.

Our study seeks to address this gap by focusing on EEG-derived symmetry features, specifically within the temporal lobes. We hypothesize that symmetry and asymmetry patterns in this region capture key neurophysiological adaptations associated with antidepressant response. This hypothesis is grounded in prior evidence showing that disruptions in temporal lobe connectivity correlate with altered emotional regulation (10–12) and is further supported by findings that indicate distinct connectivity dynamics between treatment responders and non-responders (22). By leveraging graphbased analyses and machine learning, we systematically evaluate whether temporal-lobe symmetry change scores capture response-related neurophysiological signal and yield reproducible above-chance separability under strict out-of-sample validation.

Understanding these symmetry aspects may help refine neurophysiological models of early treatment-related network reorganization in depression and motivates the search for objective EEG markers that can be evaluated in predictive contexts. By focusing on regions that show meaningful symmetry patterns, our work aims to provide interpretable candidate features that can be tested in larger and more heterogeneous cohorts and, if validated, integrated with other established EEG markers in future multi-marker approaches.

In this study, our primary objective is to assess whether symmetry-based EEG features from the temporal lobe reflect meaningful treatment-related neural changes in MDD patients. We use statistical analysis to quantify group-level differences between responders and non-responders. To complement these findings and evaluate their potential utility in predictive contexts, we also explore how these features perform in standard machine learning classification pipelines. However, our focus remains on interpretability and statistical robustness, rather than optimizing predictive performance. Accordingly, the predictive analysis is intended to quantify out-of-sample separability and its uncertainty under strict validation, not to propose a deployable clinical decision-support system. We therefore present the ML results as an indicator of reproducible signal that motivates external validation and integration with complementary biomarkers.

Recent work highlights both the promise and current limits of EEG-based early response prediction in MDD. In a machine learning meta-analysis of EEG studies, Watts et al. reported pooled performance of 83.93% accuracy (95% CI: 78.90–89.29) and AUC = 0.850 (95% CI: 0.747–0.890), underscoring that reliable prediction is feasible but remains imperfect across heterogeneous cohorts and pipelines (23). Because pooled meta-analytic estimates aggregate diverse outcomes and validation designs, they should be interpreted as a high-level benchmark rather than as an expected performance level for any single cohort or feature family. Moreover, time-series-aware feature representations have recently been explored. For example, a motif-discovery framework that models recurring temporal patterns in day-7 EEG achieved a cross-validated validation accuracy of 0.731 (F1 = 0.722) in the *α* band for distinguishing responders from non-responders (24). Together, these benchmarks motivate complementary, interpretable EEG representations, such as the temporal-lobe symmetry features studied here, while emphasizing the need for continued methodological development and external validation prior to clinical deployment.

## Methods

### Participants and Clinical Assessment

This study was part of a larger effort to understand and improve treatment outcomes for people living with MDD. It was approved by the ethical committee of the National Institute of Mental Health in Klecany, Czech Republic, and all procedures were conducted in line with the Declaration of Helsinki guidelines (Fortaleza, Brazil, 2013).

To assess the severity of depressive symptoms and track treatment progress, we used two widely recognized clinical scales. The Montgomery-Åsberg Depression Rating Scale (MADRS) (25) evaluates a range of symptoms such as mood, sleep, appetite, energy levels, and feelings of guilt. Additionally, the Clinical Global Impression (CGI) scale (26) offers a standardized approach to assess overall illness severity and observe improvements over time, based on the clinician’s judgment.

The participants in this study were recruited from the Open Department of the Prague Psychiatric Centre/National Institute of Mental Health. Each had been diagnosed with MDD and met specific inclusion criteria: a minimum score of 20 on the MADRS scale and at least a CGI score of 4 at the outset of the study. To determine treatment response, we defined a positive outcome as a reduction of 50% or more in the MADRS score by the end of the treatment period. Additional details on participant selection can be found in previous publications (27–29).

### EEG Recording Protocol

For our EEG analysis, we used a dataset from 188 MDD patients collected as part of a clinical trial investigating predictors of antidepressant response. EEG recordings were gathered using a 19-electrode system aligned with the international 10-20 configuration (30). Electrodes were placed to capture signals from key brain areas, frontal, central, temporal, parietal, and occipital, with specific sites at Fp1, Fp2, F3, F4, C3, C4, P3, P4, O1, O2, F7, F8, T3, T4, T5, T6, Fz, Cz, and Pz.

Each participant underwent two resting-state EEG sessions: one at baseline, before starting antidepressant treatment, and another one week into treatment. During each session, participants rested with their eyes closed for a 10-minute period, while EEG data were recorded using a Brainscope differential amplifier (M&I, Czech Republic). The reference electrode was placed at the FCz electrode site, located midline between Fz and Cz.

To ensure high-quality recordings, we kept scalp electrode impedance below 5 kΩ, with a difference of no more than 1 kΩ between homologous electrodes. EEG signals were sampled at either 250 Hz or 1000 Hz, recorded with a 16-bit depth for a fine resolution of 7.63 nV per bit, and captured within a dynamic range of ± 250 *µ*V. The data underwent digital filtering, with high-pass and low-pass cut-offs set at 0.15 Hz and 70 Hz, respectively, to optimize signal clarity. Importantly, previous findings suggest that early shifts in EEG signals (31–35), even after just one week of treatment (22, 27– 29, 34, 36–38), may predict stable responses, highlighting the potential of EEG data from this setup as an early indicator of long-term outcomes.

### Participant Data

The full EEG dataset initially included 188 patients diagnosed with MDD. After excluding recordings with technical issues, such as excessive distortion or more than three silent channels, the final sample comprised 176 participants (48 men, 128 women), aged 18-65 years. Of these, *N*_resp_ = 84 were classified as responders and *N*_nonresp_ = 92 as non-responders, according to the MADRS criterion of ≥ 50% reduction. Baseline demographic and clinical characteristics of the full cohort and of the responder/non-responder subgroups are summarized in Table 1. Figure 1 shows the age distribution for responders and non-responders. The distributions are broadly similar across the two groups, consistent with the non-significant age difference reported in Table 3, suggesting that age is unlikely to be a major confounding factor in the observed EEG-based differences or prediction results. The figure uses violin plots to visualize the data distribution, with the median age indicated by a horizontal black line.

**Table 1:** Demographic and clinical characteristics of the dataset and responder/non-responder subgroups. Values are mean ± SD or median [Q1, Q3] unless otherwise indicated. *N*, number of patients. Q1, first quartile (25th percentile); Q3, third quartile (75th percentile). Change in MADRS score was computed as baseline minus final MADRS; MADRS percentage change was computed as 100 × (baseline − final)/baseline, so that higher values indicate greater symptom improvement.

|  | All patients | Responders | Non-responders |
| --- | --- | --- | --- |
| Sample size, $N$ | 176 | 84 | 92 |
| Age (years), mean ± SD | 45.8 ± 11.3 | 47.7 ± 10.4 | 44.1 ± 11.9 |
| Age (years), median [Q1, Q3] | 48.0 [37.0, 54.2] | 50.0 [38.8, 55.2] | 46.0 [35.0, 54.0] |
| Sex = Female, $n$ (%) | 128 (72.7%) | 65 (77.4%) | 63 (68.5%) |
| Sex = Male, $n$ (%) | 48 (27.3%) | 19 (22.6%) | 29 (31.5%) |
| Baseline MADRS score, mean ± SD | 27.7 ± 4.0 | 27.1 ± 3.6 | 28.2 ± 4.2 |
| Baseline MADRS score, median [Q1, Q3] | 27.0 [25.0, 30.0] | 26.0 [25.0, 29.0] | 28.0 [25.0, 31.0] |
| Final MADRS score, mean ± SD | 16.2 ± 8.3 | 9.2 ± 3.6 | 22.7 ± 5.7 |
| Final MADRS score, median [Q1, Q3] | 16.0 [10.0, 22.0] | 10.0 [7.0, 12.0] | 22.0 [18.0, 27.0] |
| Change in MADRS score, mean ± SD | 11.5 ± 7.8 | 17.9 ± 4.4 | 5.5 ± 5.1 |
| Change in MADRS score, median [Q1, Q3] | 12.0 [6.0, 17.0] | 17.5 [15.0, 20.0] | 6.0 [2.0, 9.0] |
| MADRS percentage change, mean ± SD | 41.7 ± 28.0 | 66.1 ± 12.6 | 19.4 ± 17.5 |
| MADRS percentage change, median [Q1, Q3] | 43.3 [21.2, 63.0] | 63.8 [55.6, 73.3] | 21.8 [8.3, 32.0] |

**Table 2:** Classifiers and hyperparameter search space explored in the inner cross-validation loop (AUC objective).

| Classifier | Hyperparameter grid |
| --- | --- |
| **Logistic Regression (LR)** | $C \in \{0.1, 1, 10\}$; $\ell_1$ ratio $\in \{0.0, 0.25, 0.5, 0.75, 1.0\}$; penalty: Elastic-Net; solver: SAGA; max_iter = 5000. |
| **Support Vector Machine (SVM)** | kernel $\in \{\text{linear}, \text{rbf}\}$; $C \in \{10^{-2}, 10^{-1}, 1, 10, 100\}$; $\gamma \in \{\text{scale}, 10^{-3}, 10^{-2}, 10^{-1}\}$ (RBF only); class_weight = balanced. |
| **Random Forest (RF)** | $n_{\text{estimators}} \in \{300, 600\}$; max depth $\in \{\text{None}, 5, 10\}$; min samples leaf $\in \{1, 3, 5\}$; class_weight = balanced. |
| **Naive Bayes (NB)** | No hyperparameters; Gaussian likelihood assumption with class prior estimated from training data. |
| **Multilayer Perceptron (MLP)** | hidden layer sizes $\in \{(50), (100), (50, 50)\}$; activation $\in \{\text{relu}, \text{tanh}\}$; $\alpha \in \{10^{-4}, 10^{-3}, 10^{-2}\}$; early stopping = True; max_iter = 3000. |

**Table 3:** Comparison of demographic and clinical variables between responders and non-responders. Continuous variables are summarized as mean ± SD and were compared using Mann–Whitney U tests; sex distribution was compared using a chi-square test. Effect sizes for continuous variables are reported as Cohen’s *d*; effect size is not reported for the categorical sex variable.

| Variable | Responders | Non-responders | Test | p-value | Cohen’s $d$ |
| --- | --- | --- | --- | --- | --- |
| Age (years) | 47.7 ± 10.4 | 44.1 ± 11.9 | Mann–Whitney U | 0.0577 | 0.32 |
| Baseline MADRS score | 27.1 ± 3.6 | 28.2 ± 4.2 | Mann–Whitney U | 0.0648 | −0.29 |
| Final MADRS score | 9.2 ± 3.6 | 22.7 ± 5.7 | Mann–Whitney U | 7.71e-30 | −2.82 |
| Change in MADRS score | 17.9 ± 4.4 | 5.5 ± 5.1 | Mann–Whitney U | 3.54e-28 | 2.61 |
| MADRS percentage change | 66.1 ± 12.6 | 19.4 ± 17.5 | Mann–Whitney U | 2.51e-30 | 3.04 |
| Sex distribution (Female/Male) | 65 / 19 | 63 / 29 | Chi-square | 0.2480 | — |

**Table 4:** BY-adjusted *p*-values (*q*_BY_) for between-group Mann-Whitney U tests on Δ = V2 − V1 (region-wise and electrode-wise symmetry). Significant values (*q*_BY_ *<* 0.05) are shaded.

| Parameter | $\delta$ | $\theta$ | $\alpha$ | $\beta_1$ | $\beta_2$ | $\gamma$ |
| --- | --- | --- | --- | --- | --- | --- |
| RSTcore_bin | 1.000 | 0.7673 | 0.0015 | 0.0108 | 0.0276 | 0.2636 |
| RSTcore_wei | 1.000 | 1.0000 | 0.0045 | 0.0466 | 0.0601 | 0.4192 |
| RSText_bin | 1.000 | 1.0000 | 0.0119 | 0.0021 | 0.0695 | 1.0000 |
| RSText_wei | 1.000 | 1.0000 | 0.0161 | 0.0103 | 0.0161 | 1.0000 |
| ES_T3/4_bin | 1.000 | 0.5406 | 0.2683 | 0.0096 | 0.1744 | 1.000 |
| ES_T3/4_wei | 1.000 | 0.8699 | 0.2833 | 0.0465 | 0.0925 | 1.000 |
| ES_T5/6_bin | 1.000 | 0.9801 | 0.0147 | 0.0018 | 0.0960 | 1.000 |
| ES_T5/6_wei | 1.000 | 0.6007 | 0.0079 | 0.0079 | 0.0512 | 1.000 |

**Figure 1.**
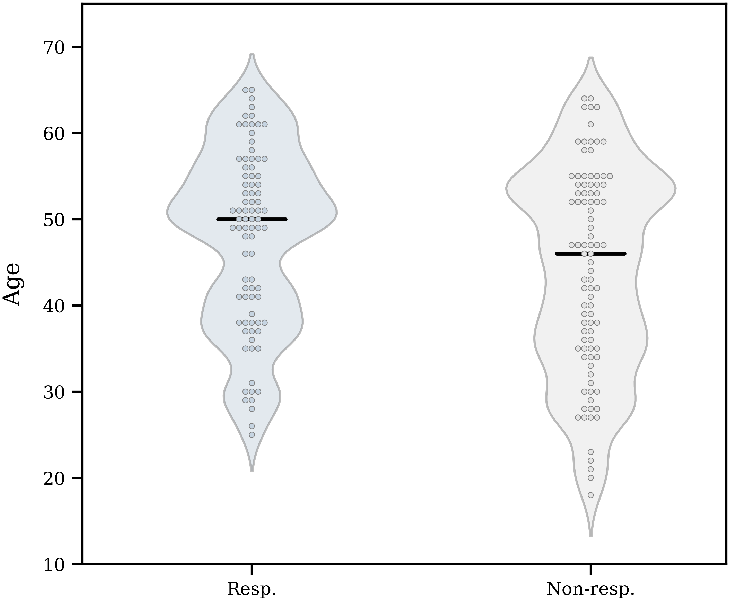
Age distribution for responders and non-responders. The violin plots display the full distribution of participant ages in each group, with individual data points overlaid and the group median indicated by a horizontal black line.

### Preprocessing and Artifact Rejection

To ensure data uniformity, the recordings that were originally sampled at 1000 Hz were downsampled to 250 Hz. In order to minimize artifacts associated with subject movement and settling, the first and last 30 seconds of each recording were discarded. Subsequently, the EEG data were re-referenced to an average reference and band-pass filtered between 1 and 40 Hz to isolate the frequency range of interest.

Following these initial steps, the artifact-reduced EEG was segmented into non-overlapping 2-second windows. Each window was assessed by evaluating the power distribution in each channel. A window was marked for exclusion if more than 20% of its channels exhibited power values that deviated significantly from a robust estimate of typical channel power. Specifically, power values were compared against thresholds derived from the median and median absolute deviation (MAD) of the channel’s power distribution, ensuring insensitivity to outliers. This method is implemented using the `clean_windows` function from the EEGLAB toolbox and effectively removes segments contaminated by brief, high-amplitude artifacts. Eight consecutive 2-second windows were then aggregated to yield 16-second segments, thereby ensuring both temporal continuity and sufficient data length for robust connectivity estimation.

### Wavelet-Based Coherence Estimation

Spectral decomposition of the EEG signals was performed using the complex Morlet wavelet, which is particularly suited for the analysis of non-stationary signals. The complex Morlet wavelet is defined as

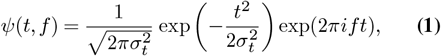

where *σ*_*t*_ determines the temporal width of the Gaussian envelope and *f* is the central frequency. The wavelet transform of a signal *X*(*t*) is then

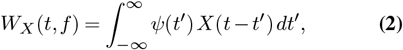

yielding complex coefficients *W*_*X*_ (*t, f*) that encode both amplitude and phase information.

For each 2-second window, wavelet coefficients were computed at 78 frequencies uniformly distributed between 1 and 40 Hz. To quantify interactions between two channels, *X*(*t*) and *Y* (*t*) during a time window, the cross-wavelet coefficients were computed as

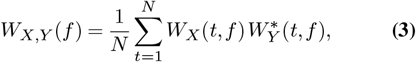

where 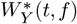 denotes the complex conjugate of *W*_*Y*_ (*t, f*) and *N* is the number of time points within the window. Coefficients from eight consecutive 2-second windows were aggregated to yield estimates over a 16-second segment.

Functional connectivity between two channels was quantified by means of the complex wavelet coherence, defined as

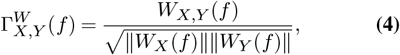

where ∥ *W*_*X*_ (*f*) ∥ is the norm of the wavelet coefficients computed over the 16-second segment. In order to suppress spurious connectivity effects due to volume conduction, the lagged coherence measure (39, 40) is considered. A refined connectivity index is therefore computed as

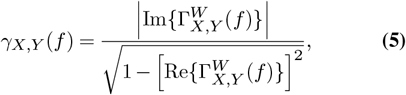

which accentuates physiologically relevant phase relationships. Note that other authors (see for example (41)) use the term *lagged coherence* to refer to the full coherence of lagged signals.

The statistical significance of the connectivity estimates was determined via surrogate data analysis. For each channel, 30 surrogate datasets were generated by randomizing the phase of the original signal while preserving its amplitude spectrum. Coherence was then recomputed for each surrogate, yielding a null distribution. For each 16-second segment, a Z-score was calculated by subtracting the mean surrogate coherence from the observed coherence and dividing by the standard deviation of the surrogate values. Connectivity estimates with Z-scores greater than 1.65 were considered statistically significant.

Subsequently, significant coherence values computed at 78 frequencies were aggregated into six typical EEG bands: *δ* (1.5 to 3.5 Hz), *θ* (4 to 7.5 Hz), *α* (8 to 12 Hz), *β*_1_ (12.5 to 17.5 Hz), *β*_2_ (18 to 25.5 Hz), and *γ* (26 to 40 Hz). Both weighted and binary connectivity matrices were constructed for these frequency bands and used to derive network measures such as cross-hemispheric connectivity and lateral asymmetry, which serve as candidate biomarkers of early treatment response.

This preprocessing and analysis pipeline, which integrates rigorous artifact removal, precise temporal segmentation, wavelet-based spectral decomposition, and robust statistical evaluation, aims for the derived functional connectivity metrics to reliably reflect underlying physiological interactions in the EEG. For further details on the entire preprocessing procedure and data characteristics, see (22).

### Graph Construction

The 19 electrodes used for EEG recording are arranged according to the international 10–20 system, as illustrated in Figure 2. Based on the preprocessing described in the previous sections, we convert the data recorded in each visit into a set of ordered graphs. Moreover, the edge set of each of the graphs is associated with weights computed through the coherence calculation process explained in Section. Therefore, the recorded data of each patient in each visit is converted into a set of ordered edge weighted graphs on 19 vertices. In fact, we constructed both weighted and binary versions of the connectivity graphs. In the weighted form, each edge retains the coherence value as its weight, thereby reflecting the strength of interaction between electrode pairs. In contrast, the binary form disregards magnitude and encodes only the presence or absence of statistically significant connectivity. In Section, we define graph-based symmetry/asymmetry features and focus on visit-to-visit change scores (Visit 2 − Visit 1) as candidate early markers of treatment response. Statistical testing and longitudinal modeling (Section) are then used to quantify within-group changes and between-group differences in these features.

### Feature Description

Herein, we provide a detailed description of the features considered in this study. Previous research (10–12) indicates that brain connectivity and functional networks exhibit symmetric properties. We focus on folding the graph along the central line containing the vertices *Fz, Cz*, and *Pz*. This line will be referred to as *the folding line* or *the central line*. Folding along this line results in a symmetric pairing of vertices across the left and right hemispheres. This symmetry facilitates both symmetric and asymmetric analyses of brain connectivity, which underpin the features defined in this section. The electrode positions and their symmetric pairings across the hemispheres, used to define the features below, are shown in Figure 2.

**Figure 2.**
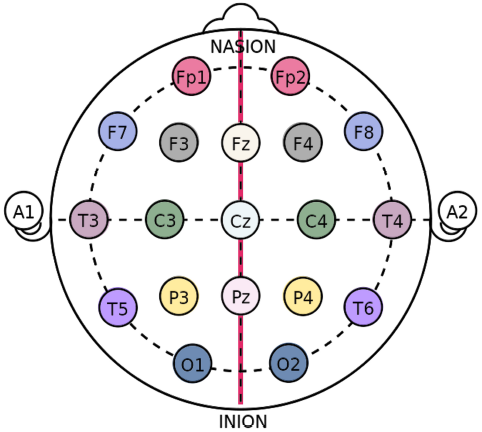
Positions of the 19 electrodes on the scalp. Electrode pairings between the left and right hemispheres are indicated using the same color. Electrodes located on the midline are considered part of both hemispheres and are paired with themselves.

This vertex pairing implicitly defines a pairing among the links, i.e., the coherences computed in Section. For instance, the coherence between channels F3 and P3 is paired with the coherence between channels F4 and P4. This link pairing allows for symmetry and asymmetry analyses of various factors, such as strength and clarity, for the paired coherences. In this paper, we consider two types of parameters: region-wise parameters and electrode-wise parameters. For the region-wise parameters, we define a subset of electrodes on the left hemisphere and analyze the paired channels on the right hemisphere relative to the folding line. Each link in the graph with both endpoints in the selected subset of electrodes on one hemisphere, along with its symmetric counterpart on the opposite side, is considered. The average weight of these paired links contributes to the parameter’s value. Formally, let *G* = (*V, E*) be the modeled graph with vertex set *V* containing 19 channels and edge set *E*. Let *S* ⊆ *V* denote the selected subset of channels on the left hemisphere, and let *E*_*S*_ := { *e* = { *u, v*} ∈ *E* : *u* ∈ *S, v* ∈ *S}* denote the set of links with both endpoints in *S*. The region-wise symmetry parameter is then given by

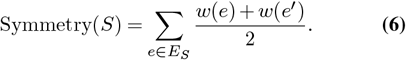

Here, *e* and *e*^*′*^ are paired (mirrored) links across hemispheres; only pairs for which both *e* and *e*^*′*^ exist in the graph are included in the sum. The function *w* : *E*(*G*) → ℝ assigns the coherence value computed in Section to each link.

For the electrode-wise parameters, we fix a channel on the left hemisphere and consider all links in the left hemisphere that have one endpoint at the selected channel. We then average the weights of paired links across both hemispheres. Specifically, if the selected channel on the left hemisphere is *v* ∈ *V* (*G*), then the parameter is defined analogous to Equation Eq. (6), where *E*_*v*_ is now the set of all links with one endpoint at *v*, and *e*^*′*^ is the paired link with *e*. Note that only pairs where both *e* and *e*^*′*^ exist are included in the sum. Additionally, a binary version of each parameter, based solely on the presence or absence of links, is also considered in this study.

During preprocessing, the recorded EEG data for each frequency band and visit is partitioned into segments, resulting in multiple weighted graphs. The computed parameter for each graph is averaged over these segments, and this average value is reported for each patient in each frequency band and visit.

We define four symmetry-based features to characterize cross-hemispheric EEG coherence. Two of these are region-wise and two are electrode-wise.

The first region-wise parameter, denoted as RSTcore, corresponds to the regional symmetry in the main temporal lobe and is defined over channels T3, T5, and O1 on the left hemisphere and their counterparts T4, T6, and O2 on the right (Figure 3a). Specifically, for each connection T3-T5, T3-O1, and T5-O1, the coherence value is averaged with its mirrored counterpart (T4-T6, T4-O2, or T6-O2), provided that both links exist, as defined in Equation Eq. (6). In the binary version of the parameter, a mirrored pair contributes a value of 1 if both links are present in the graph, and 0 otherwise, irrespective of their coherence values.

**Figure 3.**
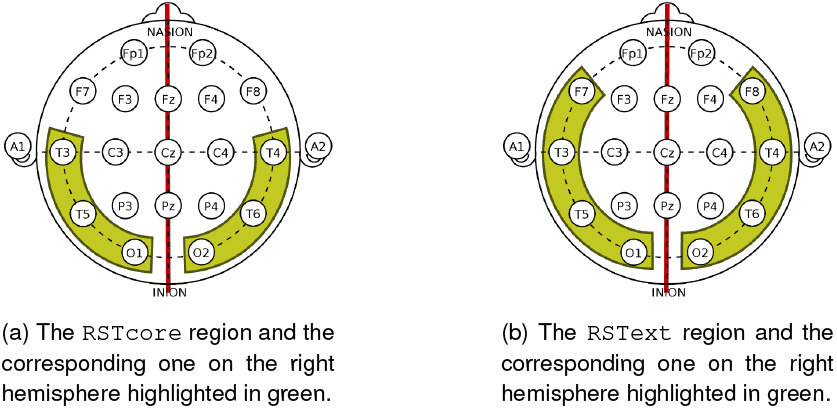
Region-wise parameters on both hemispheres. In both cases, the folding line is highlighted in red.

**Figure 4.**
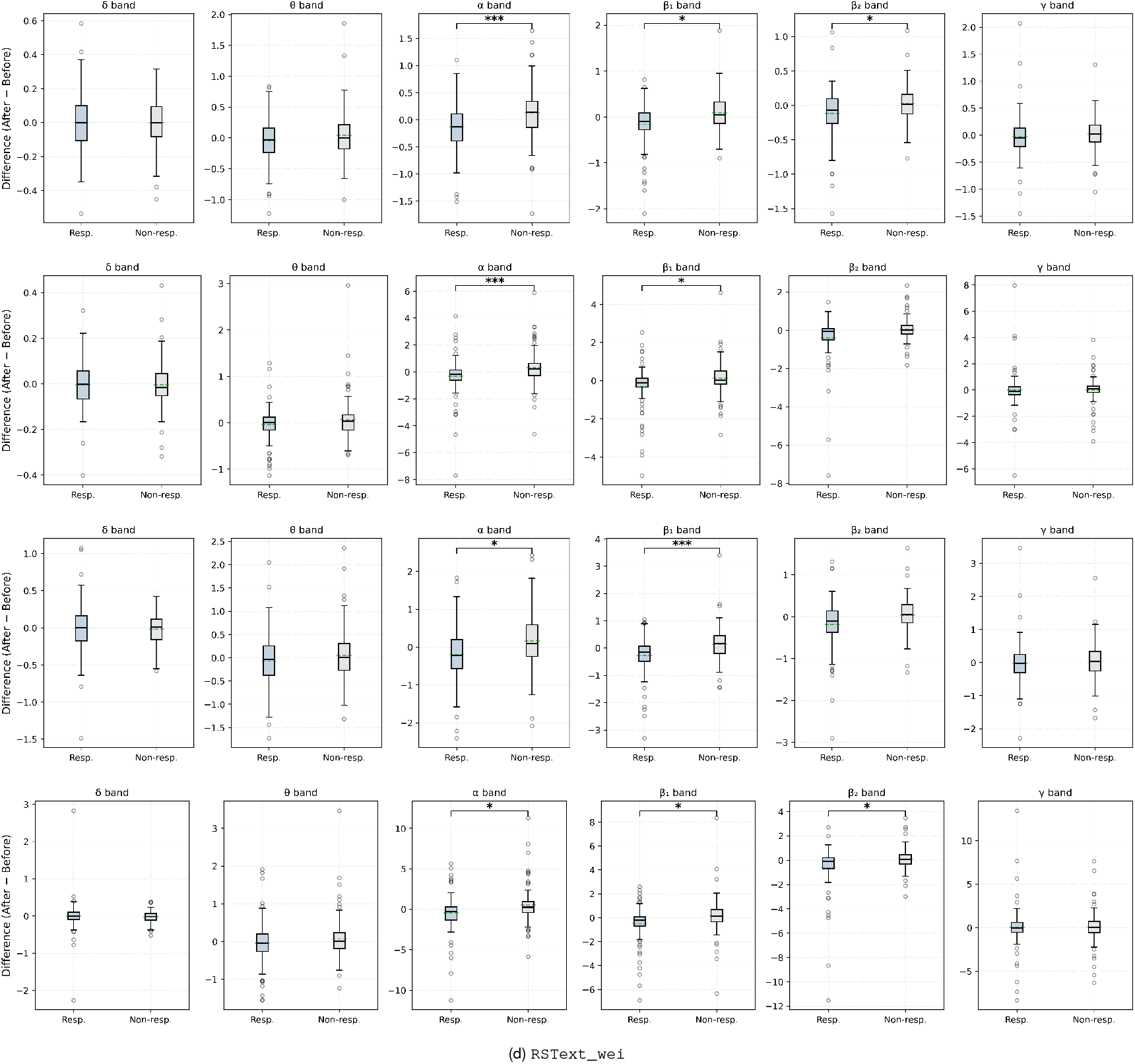
Distribution of before-after changes (Δ = V2 − V1) for region-wise symmetry parameters across six frequency bands. Responders and non-responders are shown side-by-side. Panels are annotated with BY-corrected Mann-Whitney U *q*_BY_ for the *between-group* Δ contrast: * *q <* 0.05, ** *q <* 0.01, *** *q <* 0.005.

**Figure 5.**
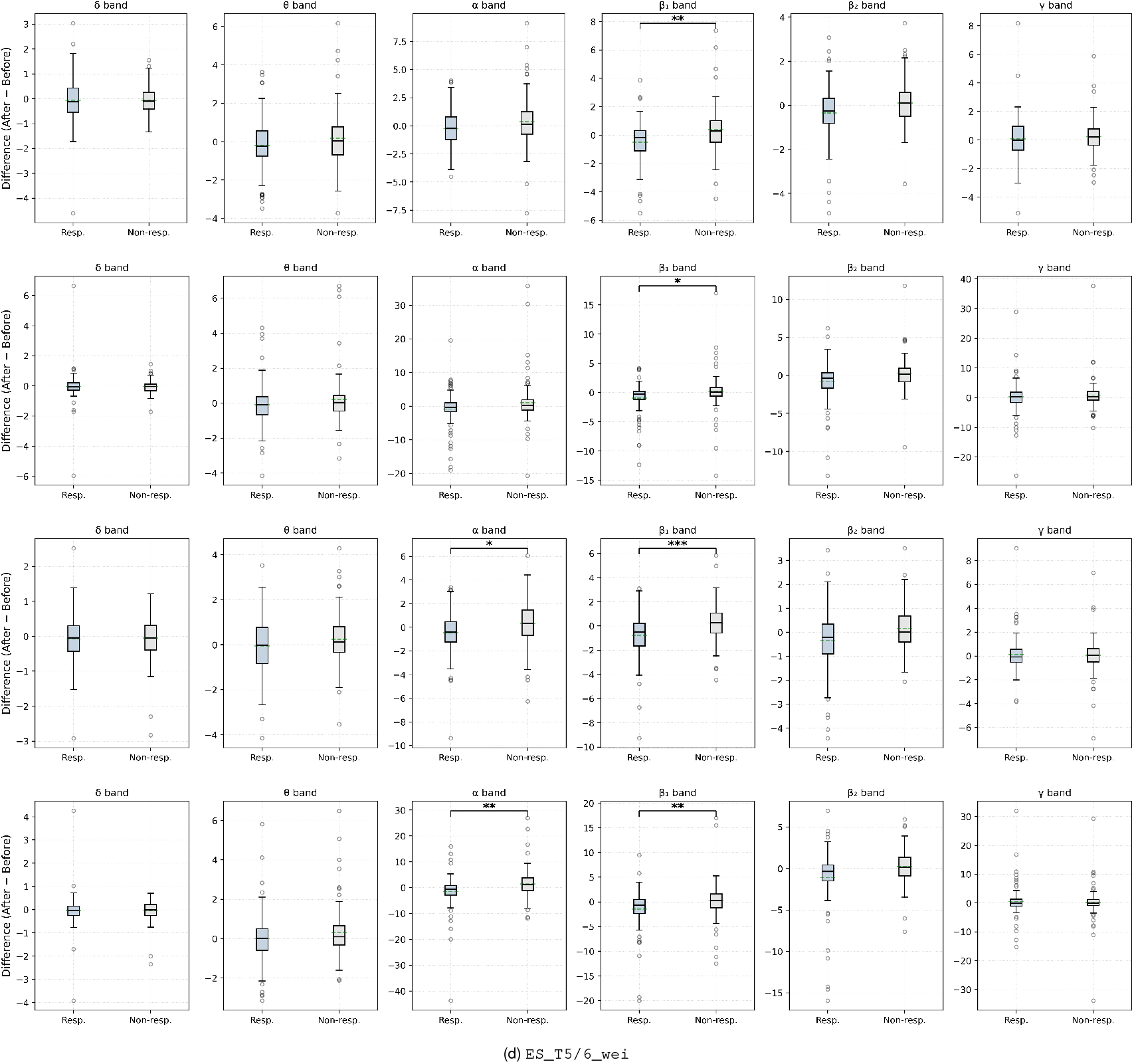
Distribution of before-after changes (Δ = V2 − V1) for electrode-wise symmetry parameters (T3-T4 and T5-T6). Responders and non-responders are shown side-by-side with BY-corrected Mann-Whitney U markers for the between-group Δ bootstrap confidence interval across outer folds of the repeated nested cross-validation. Results highlight the reproducible discriminative power of the α and *β*_1_ bands, while lower and higher frequency bands perform near chance level. For each band, the feature selector and classifier achieving the highest AUC are shown.

The second parameter, RSText, extends this region to include F7 on the left and F8 on the right (Figure 3b), thereby covering T3, T5, O1, F7 and their mirrored counterparts.

These two temporal masks are used as a robustness and spatial-extent check, that is, to examine whether the symmetry phenomenon is confined to a focal temporal cluster (RSTcore) or distributed across a slightly broader fronto– temporal region (RSText), rather than to posit qualitatively distinct neural mechanisms.

Additionally, we define two electrode-wise features of symmetry focused on the temporal lobe: ES_T3/4 and ES_T5/6. Each feature compares the set of edges connected to a given electrode in the left hemisphere (e.g., T3) with the mirrored set on the right (e.g., T4), averaging the paired edge values across hemispheres. Both binary and weighted versions of each feature are again considered.

These features were explicitly defined within the temporal lobe based on prior evidence linking this region to emotional regulation and MDD response. Thus, all subsequent analyses are focused specifically on symmetry within this region, rather than a whole-brain exploratory approach.

### Statistical Tests

The statistical analysis was designed to disentangle treatment-induced changes from pre-existing group differences and to evaluate whether responders and non-responders followed distinct temporal trajectories. All tests were conducted separately for each network parameter (defined in Section) and frequency band, using nonparametric methods that make minimal distributional assumptions and remain robust to potential outliers or skewness typical of EEG-derived measures. The overall framework consisted of two complementary steps: (i) a nonparametric comparison framework based on the Wilcoxon signed-rank and Mann-Whitney U tests, and (ii) a model-based confirmation using longitudinal generalized estimating equations (GEE).

We first examined baseline (Visit 1) values to ensure that responders and non-responders did not differ before treatment initiation. Demonstrating baseline comparability is essential for attributing any subsequent divergence to treatment effects rather than to pre-existing group differences. These baseline contrasts were assessed with two-sided Mann-Whitney U tests, applied independently to each parameter and frequency band.

Next, we quantified treatment-related change by computing, for each subject, a difference score defined as Δ = Visit 2 − Visit 1. This difference captures the early impact of treatment and provides an interpretable measure of within-subject change. Two complementary statistical questions were then addressed.

First, we assessed whether each group exhibited a significant within-subject change between visits. This was tested using the Wilcoxon signed-rank test, separately for responders and for non-responders, across all features and bands. This step identifies which parameters showed a systematic shift after treatment within each cohort, thereby indicating whether measurable neural changes occurred even in the absence of between-group divergence.

Second, we examined whether the magnitude of change itself differed between groups. This was done by applying the Mann-Whitney U test to the distributions of Δ values for responders and non-responders. This between-group comparison focuses on the central scientific question: whether responders exhibit a statistically distinguishable pattern of change compared to non-responders. The combination of these two tests, within-group Wilcoxon and between-group Mann-Whitney, allows both the directionality and the specificity of treatment effects to be examined. Together, they provide a richer picture of how neurophysiological features evolve in response to treatment.

Recognizing that the second visit might already reflect early markers of treatment response, we also compared the raw Visit 2 values between groups using Mann-Whitney tests. This complementary analysis explores whether post-treatment differences are already evident at this early stage, potentially serving as predictive indicators of later treatment efficacy.

Given the number of features and the interdependence between frequency bands, multiple testing correction was necessary to control the risk of Type I errors. To this end, all families of *p*-values within a given analysis were adjusted using the Benjamini-Yekutieli (BY) procedure (42), which controls the false discovery rate (FDR) under arbitrary dependence among tests. The BY method is more conservative than the classical Benjamini-Hochberg procedure and is therefore particularly suited to our data, where substantial correlations between spectral bands and derived network parameters are expected. Adjusted *p*-values (*q*-values) below 0.05 were considered statistically significant. For visualization clarity, only BY-significant Δ-based comparisons were displayed in the main figures, whereas baseline and Visit 2-only results were summarized comprehensively in the tables.

To further corroborate the results obtained from rank-based tests and to explicitly account for the repeated-measures nature of the data, we fitted a set of generalized estimating equations (GEE). In these models, each parameter value was expressed as a function of group (responder vs. non-responder), time (Visit 1 vs. Visit 2), and their interaction. The group-by-time interaction term captures whether the evolution across visits differs systematically between groups, thus aligning conceptually with the Δ-based nonparametric test. GEE analyses were performed separately for each parameter and frequency band, assuming an exchangeable within-subject correlation structure and using robust (sandwich) standard errors to ensure valid inference under mild model misspecification. As a sensitivity analysis, analogous linear mixed-effects models with subject-specific random intercepts were also fitted, yielding comparable conclusions. The resulting *p*-values were again adjusted using the BY procedure to maintain control over the FDR across the six frequency bands.

This combined approach of nonparametric paired and independent rank tests supplemented by longitudinal modeling offers a statistically robust and adaptable framework. The Wilcoxon and Mann-Whitney tests provide robust, distribution-free inference on both within- and between-group effects, while the GEE framework confirms that these findings remain stable when within-subject correlation is modeled explicitly. The BY correction ensures that the probability of false positives remains controlled even under correlated comparisons, maintaining the integrity of statistical inference. Together, these layers yield a comprehensive and reliable evaluation of early treatment-related changes in network parameters, balancing sensitivity to genuine effects with stringent control of Type I error. All analyses were implemented in Python using `SciPy` and `statsmodels`.

### Machine Learning Analysis

A dedicated machine learning (ML) framework was developed to evaluate the discriminative potential of the proposed EEG symmetry measures under statistically rigorous conditions. The objective of this phase was to determine whether the extracted features contain reproducible predictive signal while avoiding any form of overfitting or information leakage. Rather than aiming to maximize classification accuracy, the analysis was designed to provide a reliable, uncertainty-aware characterization of feature behavior across multiple classifiers, frequency bands, and feature-selection strategies.

Given the cohort size and the use of subject-level, low-dimensional tabular predictors (symmetry change scores), we prioritize model families that can be trained and validated robustly under resampling while remaining interpretable. End-to-end deep learning or sequence models on raw EEG typically require substantially larger datasets and introduce additional design choices (e.g., architecture depth, temporal augmentation, and regularization) that can inflate variance and reduce transparency. Our goal here is therefore to benchmark whether the proposed symmetry features carry discriminative signal that generalizes under stringent, leakage-free evaluation across multiple standard classifier families.

For each participant, the input representation consisted of the difference between the two recording sessions (*Visit 2* − *Visit 1*), thereby capturing treatment-related changes in connectivity. Features were computed within six standard frequency bands (*δ, θ, α, β*_1_, *β*_2_, and *γ*), and an additional combined condition (*α*+*β*_1_+*β*_2_) in which the three most clinically relevant bands were concatenated to test whether pooled information improved separability. Within each band, both binary and weighted forms of four symmetry indices were retained: the region-based measures `RSTcore` and `RS-Text`, and the electrode-pair measures `ES_T3/4` and `ES_-T5/6` (see Section for precise definitions). This configuration yielded eight features per single-band condition and twenty-four features for the combined *α*+*β*_1_+*β*_2_ condition. All features were computed at the subject level and centered on visit-to-visit differences, ensuring that every predictor reflected longitudinal treatment-related changes.

To obtain unbiased generalization estimates, a strict nested cross-validation (CV) procedure was employed. An outer loop with five stratified folds was used to evaluate model performance on unseen data, while an inner loop with five folds was dedicated to hyperparameter optimization and feature selection. The entire nested procedure was repeated five times using independent random partitions. Repeating the full CV mitigates variability due to fold assignment and yields a more stable distribution of performance estimates. All preprocessing, feature selection, and parameter tuning were performed strictly within the inner loop, guaranteeing that no information from the outer-test partitions influenced model training. Nested CV was chosen because it provides an unbiased estimate of out-of-sample performance even when multiple model families and hyperparameter combinations are compared simultaneously, a setting in which ordinary *K*-fold CV is known to produce optimistic bias.

Three complementary feature-selection methods were implemented to capture distinct philosophies of variable ranking. First, `SelectKBest` performed univariate filtering using the ANOVA *f* -statistic, retaining the *k* features with the highest marginal relevance. Second, `Recursive Feature Elimination` (RFE) used a logistic regression estimator with Elastic-Net regularization to iteratively prune features according to coefficient magnitude, thereby accounting for inter-feature dependencies. Third, an embedded coefficient-based method (`TopKFromModel`) employed the same Elastic-Net model to rank features directly after a single fit. Elastic-Net regularization was selected because it combines the sparsity of *ℓ*_1_ (LASSO) with the stability of *ℓ*_2_ (ridge) penalties, reducing the influence of multicollinearity while maintaining automatic feature selection. Subset sizes were explored over *k* ∈{ 2, 4, 6, 8, 12, 24}, automatically truncated to the number of features available for each band or pooled condition. Coefficient-based selectors were preceded by standardization to ensure scale invariance, whereas for other selectors scaling was applied only after feature selection when required by the downstream classifier. This conditional sequencing ensures that feature scaling influences model coefficients without biasing feature selection.

Five representative classifiers were evaluated to span the principal methodological families used in biomedical data analysis: logistic regression (LR) with an Elastic-Net penalty (43), support vector machine (SVM) (44) with linear and radial-basis-function kernels, random forest (RF) (45), naive Bayes (NB) (46), and multilayer perceptron (MLP) (47). These classifiers were chosen to encompass linear, kernel-based, ensemble, probabilistic, and neural architectures, thereby testing whether observed effects were model-specific or intrinsic to the features themselves. Class weights were balanced where supported to mitigate potential class-imbalance bias. Hyperparameters were tuned within the inner CV loop by exhaustive grid search using the area under the receiver operating characteristic curve (AUC) as the sole optimization criterion. Using AUC rather than accuracy avoids dependence on arbitrary classification thresholds and provides a threshold-independent measure of separability that is appropriate for small or imbalanced biomedical datasets. The hyperparameter search spaces explored in the inner loop are summarized in Table 2.

At each outer fold, the best configuration identified by the inner loop, including selector type, number of selected features, and classifier hyperparameters, was refitted on the full training partition and evaluated on the corresponding test fold. This procedure was executed independently for each frequency-band and combined-band condition, across all feature selectors and classifiers, and repeated over the five randomized nested CV runs. Predicted labels, decision scores, and chosen hyperparameters were stored per outer fold; feature-selection stability was summarized via selection frequencies across folds, repeats, selectors, and classifiers. All computations were parallelized across CPU cores, and random seeds were fixed per repeat and fold to ensure deterministic reproducibility.

Statistical uncertainty was quantified using non-parametric bootstrap resampling of outer-test results. For each metric, 2000 bootstrap samples were drawn with replacement from the set of outer-fold scores, the median performance was computed within each sample, and the 2.5th and 97.5th percentiles of these medians were reported as a 95% confidence interval. Bootstrap CIs were preferred to parametric estimates because cross-validated metrics are not normally distributed, and resampling provides a distribution-free estimate of uncertainty around the median. Feature-selection stability was assessed by counting the frequency with which each feature was retained across all folds, repeats, selectors, and classifiers, thereby identifying consistently informative features irrespective of model choice or data partitioning.

The complete workflow of the analysis is summarized in Algorithm 1, which outlines the nested cross-validation, inner-loop optimization, and outer-loop evaluation sequence. Optional routines for permutation testing and learning-curve estimation were implemented but disabled in the main analyses for computational efficiency. All analyses were conducted in Python 3.11 using `scikit-learn` (version 1.5). All code, scripts, and parameter settings are openly available in the project repository.

#### Algorithm 1

Repeated Nested Cross-Validation with Integrated Feature Selection and Bootstrap Uncertainty Estimation

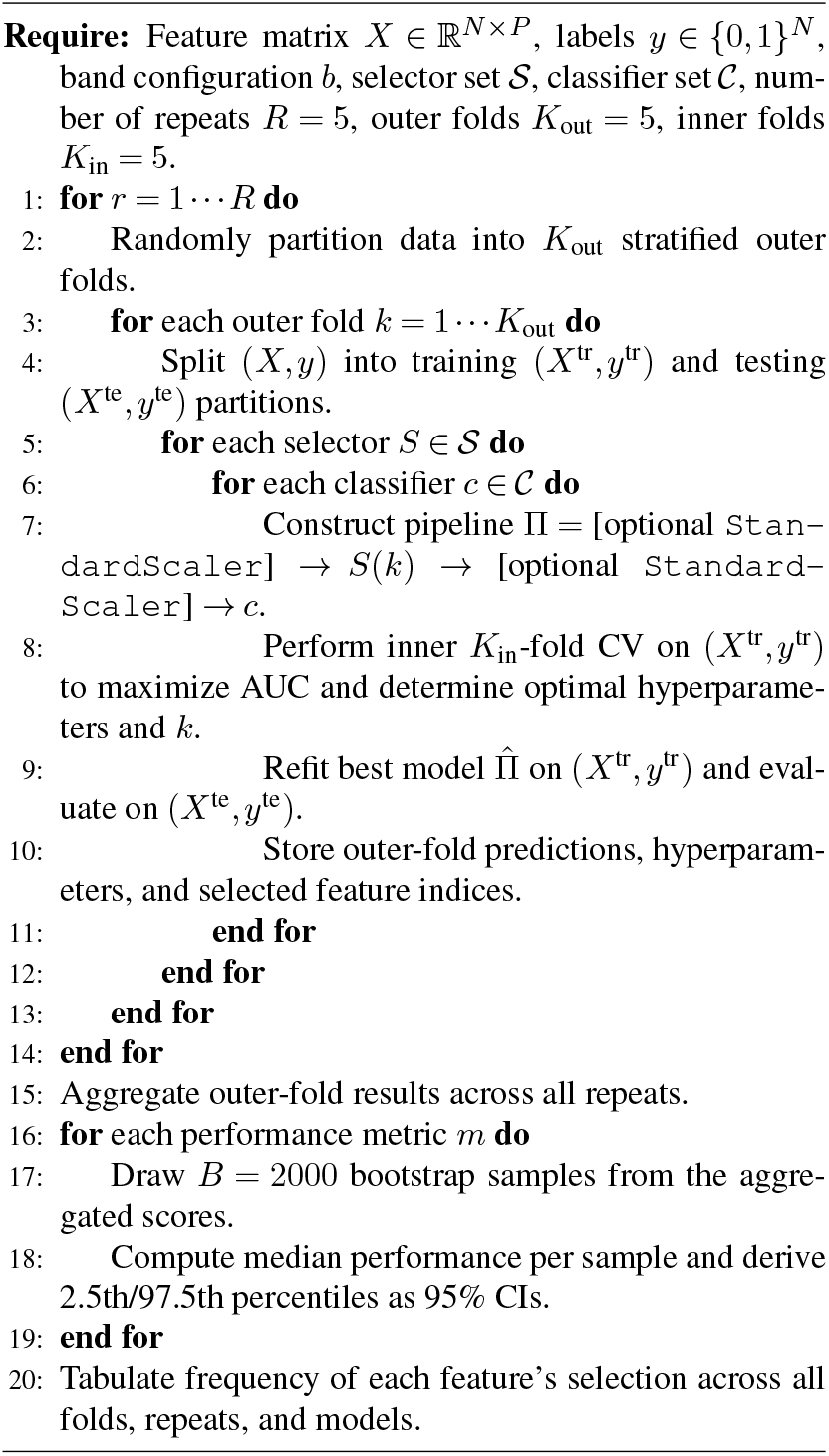

This framework provides a transparent and statistically grounded assessment of the predictive value of EEG symmetry features, incorporating robust cross-validation, selector-aware scaling, comprehensive model comparison, and non-parametric uncertainty estimation within a single reproducible pipeline.

## Results

We analyzed *N* = 176 patients (84 responders, 92 non-responders). With the cohort, preprocessing, coherence estimation, graph construction, and feature derivation set out in Sections -, this section reports the outcomes of the two pre-specified analyses described in Sections -.

Before turning to the EEG-based analyses, we examined whether responders and non-responders differed in basic clinical or demographic characteristics (Table 3). Responders tended to be slightly older than non-responders and to have marginally lower baseline MADRS scores, but these differences were small in magnitude and did not reach statistical significance (Mann–Whitney U tests for age and baseline MADRS, both *p >* 0.05; | Cohen’s *d* | ≈ 0.3). The sex distribution was also comparable between groups (*χ*^2^ = 1.33, *p* = 0.2480). As expected from the response definition, responders and non-responders differed very strongly in final MADRS scores and in absolute and percentage change (all *p* ≪ 0.001, very large effect sizes; Table 3). These comparisons indicate that large confounding of the EEG findings by age, sex, or baseline severity is unlikely.

### Statistical Results for Symmetry Features

As pre-specified in Section, we report three contrasts: baseline (Visit 1), the primary early-change contrast Δ = Visit 2 − Visit 1, and post-treatment cross-sectional (Visit 2). Multiplicity is controlled across the six bands within each feature family using Benjamini–Yekutieli (results reported as *q*_BY_); longitudinal GEE targeting the group × time interaction serves as a corroboration of the Δ-based findings.

We first examined baseline Visit 1 values for each feature and band to assess potential pre-existing differences between responders and non-responders. No feature × band comparison survived FDR control (all *q*_BY_ ≥ 0.05), and medians were comparable between groups across region-wise metrics (RSTcore, RSText; binary and weighted) and electrode-wise metrics (ES_T3/4, ES_T5/6; binary and weighted). This supports the core identification strategy, namely that any divergences observed subsequently can be attributed to treatment-related change rather than baseline imbalance. Results for these checks are summarized compactly in Appendix Table 10, which lists *q*_BY_ values for all features and bands and shows the absence of statistically reliable Visit 1 separation.

Turning to the primary estimand, we computed per-subject change scores Δ = Visit 2 − Visit 1 and tested whether the distributions of Δ differed between responders and non-responders using Mann-Whitney U tests for each feature and band, with BY correction across the six bands. The pattern was both robust and coherent: responders exhibited larger decreases (larger magnitude of Δ) than non-responders, with FDR-significant contrasts concentrated in the mid-frequency spectrum. The complete set of between-group results for all region-wise and electrode-wise features is summarized in Table 4. In total, sixteen feature × band contrasts reached *q*_BY_ *<* 0.05; eight of these fell in *β*_1_, six in *α*, and two in *β*_2_. Notably, all significant contrasts shared the same directionality, Δ_*R*_ *<* Δ_*N*_, indicating systematically greater reductions among responders. The strongest separations were concentrated at the temporal electrodes, especially T5/T6, appearing in both *α* and *β*_1_ bands and in both binary and weighted formulations (e.g., `ES_T5/6_wei`, *α*: *q*_BY_ = 0.0079, Δ_*N*_ − Δ_*R*_ = +1.8220). Additional but slightly smaller separations were observed for `ES_T3/4` in *β*_1_ in both binary and weighted forms and for the region-wise masks, primarily the weighted versions. Across the top contrasts, effects are dominated by the *α* and *β*_1_ bands. To provide concrete magnitude and direction, Table 5 lists the top contrasts with group medians of Δ and their differences; the full set of 16 significant feature–band pairs is given in Appendix Table 11.

**Table 5:**
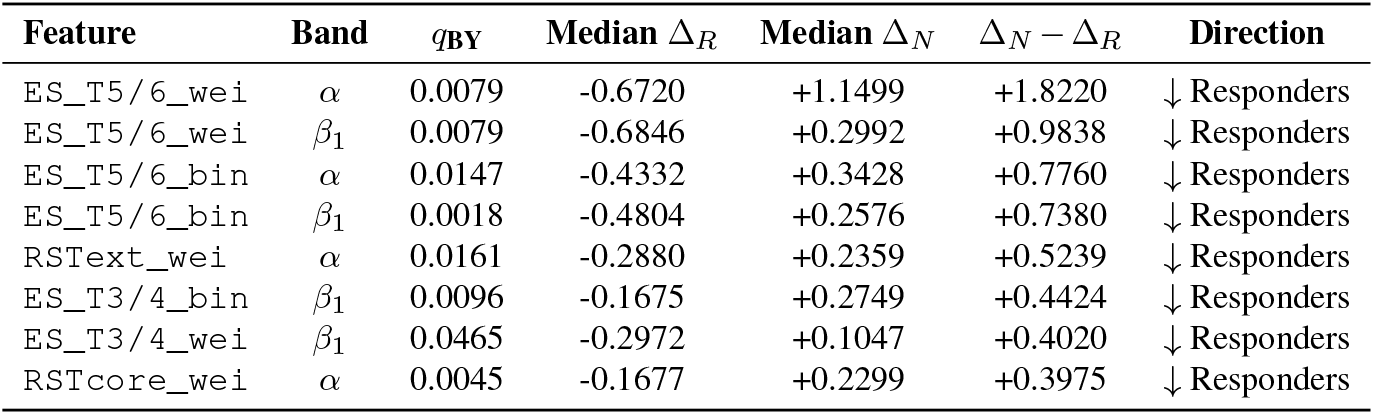
Top-8 between-group contrasts on Δ = V2 − V1 from Mann-Whitney U tests (BY-corrected across six bands per feature). Each row reports the BY-adjusted *p* (*q*_BY_), the group medians of Δ, and their difference. Negative Δ denotes a decrease from baseline. All listed contrasts indicate larger decreases in responders (↓ Responders). The full table for all the 16 significant feature-band pairs appears in Appendix Table 11.

Within-group paired Wilcoxon signed-rank tests clarified how these between-group results arise. Applied separately to responders and non-responders and BY-corrected across bands, these tests ask whether each cohort changed from baseline. Responders showed multiple BY-significant within-group decreases predominantly in *α, β*_1_ and *β*_2_ (10 shaded cells in Table 6). The strongest effects were at temporal electrodes, especially in *β*_1_: for example, `ES_T5/6_bin` (*β*_1_) had *q*_BY_ = 0.005 and `ES_T5/6_wei` (*β*_1_) had *q*_BY_ = 0.006. Region-wise masks also decreased among responders: `RSTcore_bin` in *α, β*_1_, and *β*_2_ (*q*_BY_ = 0.044 each), `RSText_bin` in *β*_1_ (*q*_BY_ = 0.016), and `RSText_wei` in *β*_1_ and *β*_2_ (*q*_BY_ = 0.022, 0.030). Additional temporal effects appeared for `ES_T3/4_wei` in *β*_1_ and *β*_2_ (*q*_BY_ = 0.019, 0.044). In contrast, no within-group shifts among non-responders survived BY correction (e.g., smallest values *q*_BY_ = 0.151-0.490 across features/bands). This triangulation is important: the between-group Δ contrast indicates differential change, while the paired Wilcoxon shows that this differential primarily reflects decreases within the responder cohort rather than increases in the non-responder cohort. Full *q*_BY_ matrices for all features and bands are provided in Table 6.

**Table 6:**
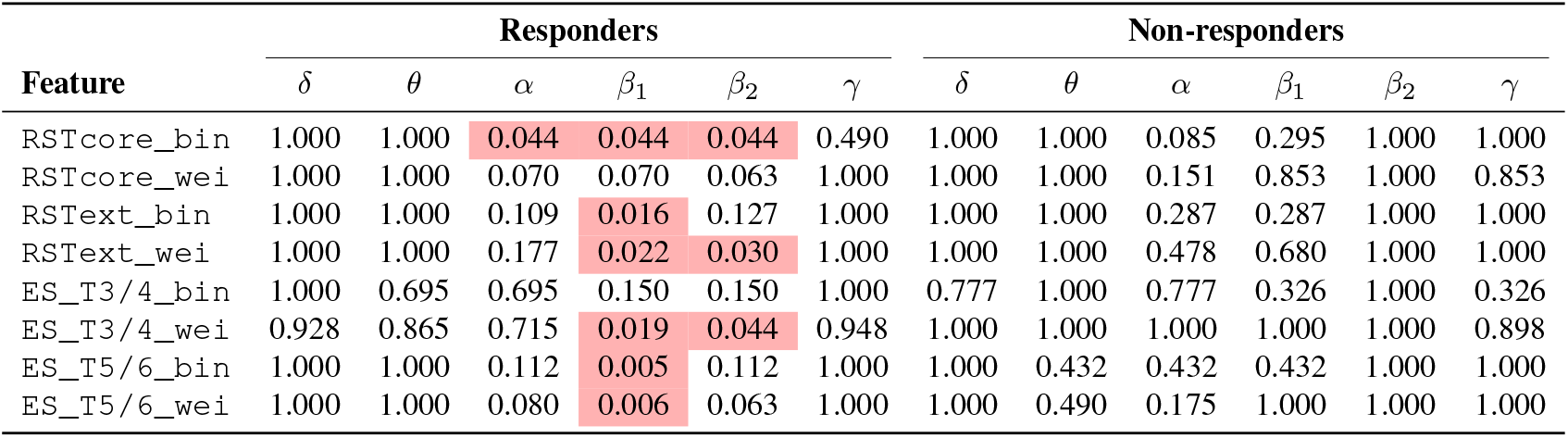
Wilcoxon signed-rank tests (Visit 2 vs. Visit 1), BY-corrected across the six bands for each feature. Left block: *responders*; right block: *non-responders*. Entries are *q*_BY_. Shaded cells denote *q*_BY_ *<* 0.05.

Because Visit 2 occurs early after treatment initiation, we also examined group differences at this time point using Mann-Whitney tests with BY correction across the six frequency bands. Several feature-band pairs (ten in total: four in *α* and six in *β*_1_) reached BY significance, all showing the same direction as the change-based results: responders exhibiting lower symmetry values than non-responders. This cross-sectional pattern indicates that measurable divergence between groups might be already emerging within the first week of treatment, consistent with the onset of early neurophysiological change. Conceptually, these single-visit differences complement rather than replace the Δ analysis: whereas Δ = V2 − V1 quantifies the difference in change between groups (the primary estimand), the Visit 2 analysis captures the earliest detectable group separation in absolute values. The overlap of significant bands between these two analyses, together with the corroborating negative group × time coefficients in the longitudinal models, supports the robustness of the early treatment effects. Detailed Visit 2 results are provided in Appendix Table 12.

To confirm that the Δ-based inferences hold under explicit modeling of within-subject correlation, we fitted GEE models for each feature and band with group, time, and their interaction; robust sandwich standard errors ensure validity under mild working-correlation misspecification. Nineteen feature × band interactions were BY-significant at *q*_BY_ *<* 0.05, and every significant coefficient was negative, indicating larger decreases over time in responders than in non-responders. The agreement with the nonparametric Δ tests was high (94%): fifteen of the sixteen FDR-significant Δ contrasts were independently corroborated by a significant group × time interaction. Also, some *β*_2_ bands reached significance in GEE that did not in the rank-based comparison, a plausible outcome given that GEE exploits the full repeated-measures structure and can recover subtler longitudinal effects when correlation is modeled. The set of GEE interactions, with coefficients and significant *q*_BY_ values, is shown in Table 7 in which *β*_interaction_ quantifies how the Visit 2 − Visit 1 change differs between responders and non-responders after modeling within-subject correlation (exchangeable working correlation; robust SEs). Thus, it targets the same scientific estimand as the between-group Δ comparison. Moreover, analogous linear mixed-effects models (random intercepts) yielded comparable conclusions (Appendix Table 13), underscoring the robustness of these effects to modeling choice.

To aid interpretation and transparency, Figures 4 and 5 visualize the Δ distributions for region-wise and electrode-wise metrics across bands, annotated with BY-corrected Mann-Whitney U results for the between-group Δ contrasts.

Forest plots of the GEE group × time coefficients with robust 95% confidence intervals (see Figure 6) convey the direction and uncertainty of longitudinal effects. Multiplicity was controlled across all feature-band tests using the BY procedure; BY-significant effects are highlighted in red. For completeness and transparency, *δ* and *γ* bands are also shown even though they exhibited no measurable interaction (BY *q*_BY_ ≈ 1 across features) and therefore appear as grey points without confidence intervals.

**Figure 6.**
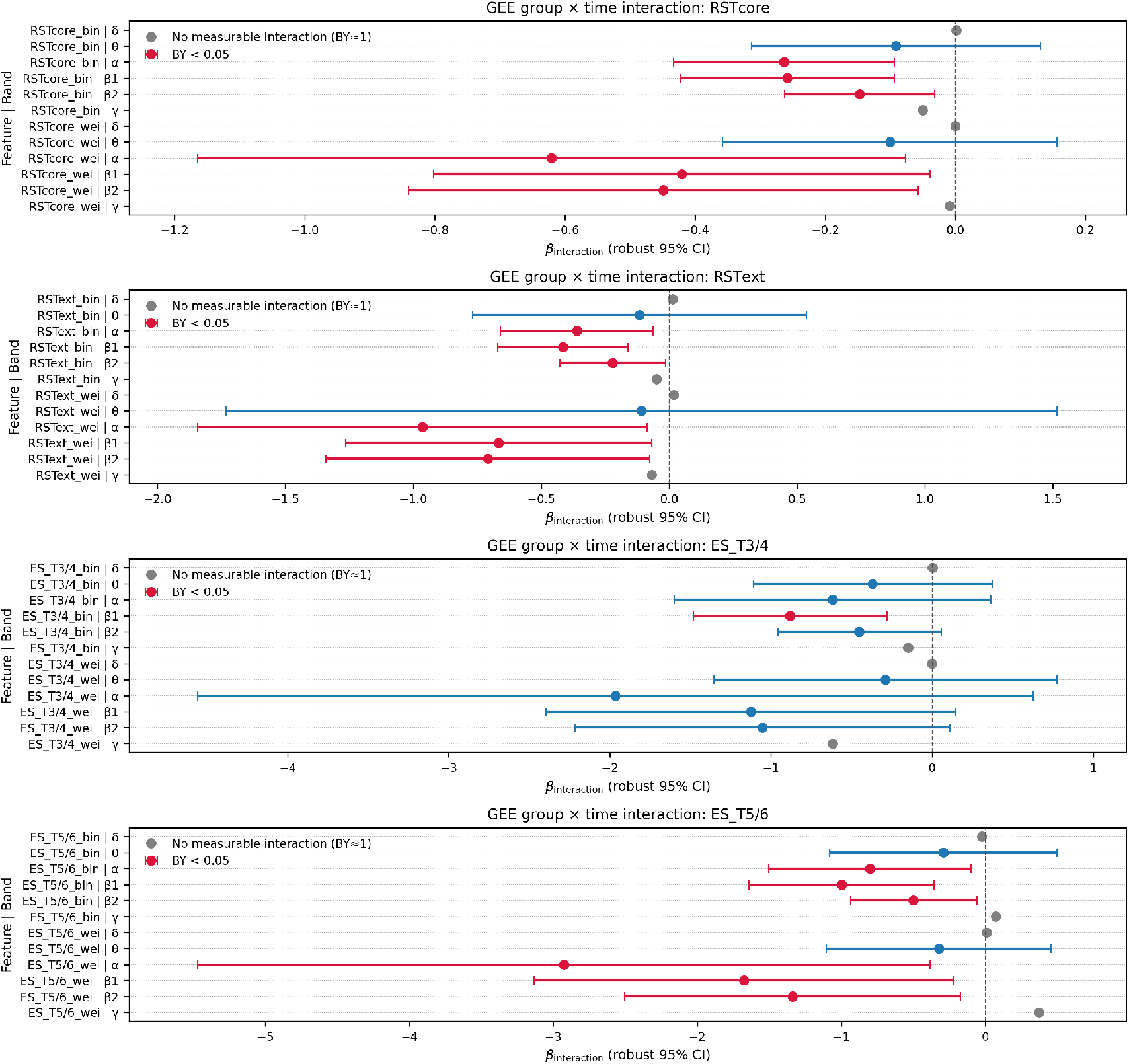
Generalized estimating equation (GEE) group × time interaction (*β*_interaction_) for each feature and the available frequency bands (*θ, α, β*_1_, *β*_2_; *δ* and *γ* included for completeness). Points show estimates with robust 95% confidence intervals where estimable; color encodes multiplicity control, red: Benjamini-Yekutieli (BY) *q*_BY_ *<* 0.05, blue: *q*_BY_ ≥ 0.05, grey: no measurable interaction (BY *q*_BY_ ≈ 1, standard error not defined). Negative coefficients indicate a larger change from Visit 1 to Visit 2 in responders relative to non-responders. Panels group features by family. All estimates are reported to emphasize effect magnitude and uncertainty; statistical significance is indicated only by color.

**Figure 7.**
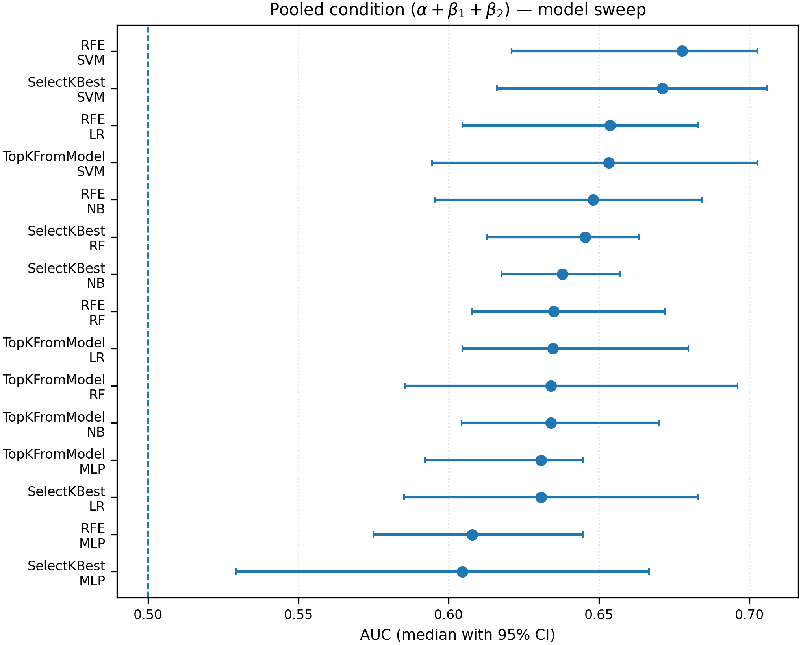
Forest plot of the median AUC and 95% bootstrap confidence intervals for all selector-classifier combinations in the pooled condition (*α*+*β*_1_+*β*_2_). Each horizontal line corresponds to one model configuration, with dots indicating the median AUC. The dashed vertical line marks chance-level performance (AUC = 0.5). The clustering of points above this baseline indicates that the separability between responders and non-responders is reproducible across different learning architectures, and confirms the stability of the discriminative single

**Table 7:** Significant GEE group × time interactions across features and bands (BY-adjusted *q*_BY_<0.05). Coefficients are from marginal models with exchangeable correlation and robust sandwich standard errors. Negative *β*_interaction_ indicates larger decreases over time in responders than in non-responders. Linear mixed-effects models yielded comparable conclusions (See appendix Table 13).

| Feature | Band | $\beta_{\text{interaction}}$ | $q_{BY}$ |
| --- | --- | --- | --- |
| RSTcore_bin | $\alpha$ | −0.264 | 0.0023 |
| RSTcore_bin | $\beta_1$ | −0.258 | 0.0021 |
| RSTcore_bin | $\beta_2$ | −0.147 | 0.0123 |
| RSTcore_wei | $\alpha$ | −0.621 | 0.0253 |
| RSTcore_wei | $\beta_1$ | −0.420 | 0.0309 |
| RSTcore_wei | $\beta_2$ | −0.449 | 0.0245 |
| RSText_bin | $\alpha$ | −0.361 | 0.0178 |
| RSText_bin | $\beta_1$ | −0.415 | 0.0013 |
| RSText_bin | $\beta_2$ | −0.221 | 0.0353 |
| RSText_wei | $\alpha$ | −0.964 | 0.0315 |
| RSText_wei | $\beta_1$ | −0.666 | 0.0290 |
| RSText_wei | $\beta_2$ | −0.708 | 0.0281 |
| ES_T3/4_bin | $\beta_1$ | −0.881 | 0.0039 |
| ES_T5/6_bin | $\alpha$ | −0.802 | 0.0255 |
| ES_T5/6_bin | $\beta_1$ | −0.998 | 0.0023 |
| ES_T5/6_bin | $\beta_2$ | −0.499 | 0.0255 |
| ES_T5/6_wei | $\alpha$ | −2.926 | 0.0241 |
| ES_T5/6_wei | $\beta_1$ | −1.676 | 0.0241 |
| ES_T5/6_wei | $\beta_2$ | −1.340 | 0.0241 |

Collectively, the evidence paints a consistent picture that is both statistically and neurophysiologically coherent: responders exhibit early reductions in interhemispheric symmetry that are most pronounced in *α* and *β*_1_, with supportive signals in *β*_2_ when within-subject correlation is explicitly modeled. The absence of Visit 1 differences reduces concern about baseline confounding; the concordance of directions across Δ medians, within-group Wilcoxon results, and GEE coefficients reinforces the interpretation that these changes are treatment-related; and the strict FDR control via Benjamini-Yekutieli across dependent bands maintains a conservative Type I error profile consistent with the correlation structure of spectral features.

### B. Classification Results

The nested cross-validation analysis provided a comprehensive assessment of the predictive value of the proposed EEG symmetry features across frequency bands, feature-selection strategies, and classifier types. As outlined in Section, the objective was not to maximize raw classification accuracy, but to determine which frequency bands carry the most reproducible predictive information and whether the resulting pattern of machine learning (ML) performance supports the statistical findings presented in Section A. All results refer to out-of-sample performance estimated from the outer folds of the repeated nested cross-validation, with uncertainty quantified as non-parametric 95% bootstrap confidence intervals of the median. Across all configurations, the classification models achieved moderate but reproducible separability between responders and non-responders. The performance distribution was narrow across folds and repeats, reflecting stable generalization rather than overfitting. The area under the receiver operating characteristic curve (AUC) was used as the primary performance metric, as it provides a threshold-independent measure of separability that is robust to class imbalance and therefore more informative than raw accuracy in biomedical datasets with moderate sample sizes.

The strongest predictive signal was consistently found in the *α* and *β*_1_ bands. The best configuration, combining recursive feature elimination (RFE) with a neural network classifier in the *α* band, achieved an outer-fold median AUC of 0.690 with a 95% bootstrap confidence interval [0.611, 0.711]. Notably, the top-performing configuration already used a non-linear neural architecture (MLP), yet the overall discrimination remained modest, consistent with a limited effect size in this cohort. A nearly identical result was observed in the *β*_1_ band when using the embedded Elastic-Net selector (`TopKFromModel`) with the same classifier (AUC_median_ = 0.686 [0.631, 0.707]). Pooling the three clinically relevant frequency bands (*α*+*β*_1_+*β*_2_) yielded com-parable but not superior performance (AUC_median_ = 0.678 [0.621, 0.703]), indicating that the discriminative information is already concentrated within the *α* and *β*_1_ ranges. The *β*_2_ band showed a weak but slightly above-chance signal (AUC ≈ 0.58), whereas *δ, θ*, and *γ* bands were indistinguishable from random performance (AUC ≈ 0.52).

Varying the number of selected features (more precisely, changing *k* over the values { 2, 4, 6, 8, 12, 24}) had only a minor influence on classification outcomes. Intermediate subset sizes (*k* = 4-8) produced the most stable results, while larger subsets occasionally introduced redundant features without improving generalization. Across all bands, both linear and non-linear classifiers (logistic regression with Elastic-Net regularization, SVM, random forest, and neural networks) produced consistent AUCs in the same frequency ranges, reinforcing that the observed effect is intrinsic to the features rather than specific to a particular modeling architecture.

Table 8 summarizes the best-performing configuration for each frequency band, reporting the median and 95% confidence interval of the outer-fold AUC. These results clearly identify *α* and *β*_1_ as the most informative frequency ranges, with confidence intervals well above the chance level, that are in line with the statistical analyses presented in Section A. This convergence between inferential statistics and predictive modeling suggests that both analyses may capture the same underlying physiological phenomenon.

The reproducibility of these findings across multiple classifiers and selectors highlights that the observed discriminative signal is not dependent on a specific modeling choice but is intrinsic to the features themselves. For example, above-chance AUCs were also obtained with simpler models such as Elastic-Net logistic regression and SVMs, indicating that the discriminative signal is not confined to a single modeling architecture (Table 8).

**Table 8.**
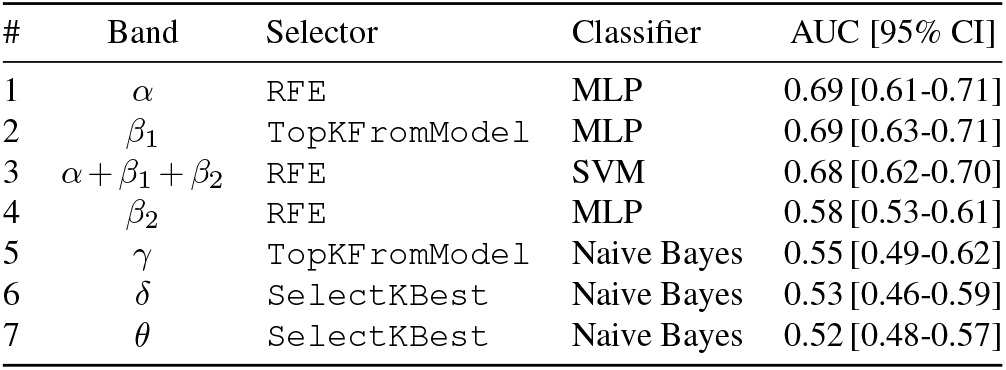
Best-performing configuration per frequency band, reporting the median AUC and its 95% Δ contrast; * *q* < 0.05, ** *q* < 0.01, *** *q* < 0.005.

| # | Band | Selector | Classifier | AUC [95% CI] |
| --- | --- | --- | --- | --- |
| 1 | $\alpha$ | RFE | MLP | 0.69 [0.61-0.71] |
| 2 | $\beta_1$ | TopKFromModel | MLP | 0.69 [0.63-0.71] |
| 3 | $\alpha + \beta_1 + \beta_2$ | RFE | SVM | 0.68 [0.62-0.70] |
| 4 | $\beta_2$ | RFE | MLP | 0.58 [0.53-0.61] |
| 5 | $\gamma$ | TopKFromModel | Naive Bayes | 0.55 [0.49-0.62] |
| 6 | $\delta$ | SelectKBest | Naive Bayes | 0.53 [0.46-0.59] |
| 7 | $\theta$ | SelectKBest | Naive Bayes | 0.52 [0.48-0.57] |

To illustrate model robustness, Figure 7 presents a forest plot of AUC medians and confidence intervals for all selector-classifier combinations in the pooled (*α*+*β*_1_+*β*_2_) condition. Each point represents an independent cross-validation configuration. The narrow confidence intervals and their clustering above the random-performance baseline (AUC = 0.5) confirm that the separability between responders and non-responders is consistent across model families and feature selection strategies, even when no single combination dominates by a large margin.

Feature-level stability analysis (Table 9) showed that the same temporal-lobe symmetry features, particularly `RSTcore_bin` in *α*, and `ES_T3/4_bin` and `ES_- T5/6_bin` in *β*_1_, were consistently retained across folds, feature selectors, and classifiers. This convergence might also underscore the physiological interpretability of the machine learning results: the same temporal-lobe symmetry features that showed the strongest group-level statistical differences also drive the predictive signal in the classification framework.

**Table 9.** Top 10 most frequently selected features across all frequency bands, feature selectors, and classifiers. Feature frequencies are aggregated across outer folds and repetitions of the nested cross-validation.

| # | Band | Feature | Selection frequency |
| --- | --- | --- | --- |
| 1 | $\alpha$ | RSTcore_bin | 95.7% (718) |
| 2 | $\beta_1$ | ES_T3/4_bin | 83.3% (625) |
| 3 | $\alpha$ | ES_T5/6_wei | 76% (570) |
| 4 | $\beta_1$ | ES_T5/6_bin | 70.1% (526) |
| 5 | $\beta_1$ | RSText_bin | 69.7% (523) |
| 6 | $\beta_2$ | RSTcore_wei | 68.9% (517) |
| 7 | $\beta_1$ | RSTcore_bin | 67.1% (503) |
| 8 | $\beta_2$ | RSText_wei | 61.3% (460) |
| 9 | $\beta_2$ | RSText_bin | 60.7% (455) |
| 10 | $\beta_2$ | ES_T5/6_bin | 58.5% (439) |

Overall, the classification results confirm that EEG symmetry features encode a stable and physiologically interpretable distinction between responders and non-responders. Although absolute classification performance remains moderate, the convergence between statistical significance and reproducible ML separability strongly supports the hypothesis that treatment-related information resides primarily in the *α* and *β*_1_ frequency bands. The consistency across model types, frequency bands, and selection strategies suggests that the predictive information is robust to modeling assumptions and likely reflects genuine neurophysiological differences rather than statistical artifacts.

## Discussion

This section places our results in context, beginning with a summary of the key statistical findings and their methodological rationale. We then interpret the symmetry changes from a neurophysiological perspective, consider how response granularity and demographic factors may refine predictive models, and outline clinical as well as research implications. Limitations and directions for future work are discussed throughout.

### C. Principal findings

Our statistical analysis framework encompassed three types of comparisons: baseline (Visit 1) differences, change between visits, and follow-up (Visit 2) differences. Among these, the change-based analysis, computed as the difference between the early-treatment and pre-treatment recordings, yielded the most consistent and significant group-level effects. The lack of significance at baseline suggests that the symmetry features are not pre-existing biomarkers of treatment response. Rather, they appear to capture early neurophysiological adaptations to treatment. Although some significant differences emerged at Visit 2 alone, these may represent the early onset of treatment-related changes that could become more pronounced over time. Future studies might incorporate additional EEG sessions later in the treatment period to track whether these asymmetry patterns continue to evolve, stabilize, or reverse. Exploring the temporal trajectory of these effects would shed light on whether the early differences observed at Visit 2 are predictive of long-term neurophysiological adaptation.

The clearest and most robust patterns in our study emerged when evaluating changes between visits. This is not only methodologically sound, but also clinically aligned with how treatment response was defined: a patient was classified as a responder only if their MADRS score was reduced by at least 50% by the end of the treatment period. Summarizing all the obtained results, three practical rules emerge. First, clinical signal lives almost entirely in the *α* and *β* bands, with *β*_1_ delivering the smallest p-values. Second, coding connectivity as present/absent amplifies the group contrast relative to weight-based formulations, suggesting that early treatment might selectively prune or add connections rather than merely modulating their strength. An additional (methodological) contributor is that binary graphs derived from statistical significance thresholds may be less sensitive to between-subject differences in signal-to-noise ratio and coherence estimation variance, whereas small weight fluctuations can dilute group contrasts. Future work should therefore explicitly quantify whether edge density and edge turnover (appearance/disappearance of significant edges) differ between groups and whether these network-level summaries mediate the observed symmetry effects. Third, whether one uses the ‘core’ or ‘extended’ temporal mask has a negligible impact, indicating the phenomenon is distributed across the lobe. Thus, evaluating EEG-based feature differences across visits provides a direct neural correlate of the clinical criteria used to define treatment success.

### D. Neurophysiological interpretation of symmetry changes

Because the present study is based on restingstate scalp EEG and does not include behavioral or task-based measures, mechanistic interpretations remain inferential. We therefore interpret the observed symmetry changes as network-level correlates of early treatment response and frame the following explanations as hypotheses to be tested in future work.

A detailed understanding of our symmetry findings requires separating (1) the mechanisms driving decreased symmetry in responders, (2) the factors underlying increased symmetry in non-responders, and (3) the special role played by temporal-lobe circuitry. We address each in turn below.

#### D.1. Decrease in symmetry among responders

The decline in symmetry observed among responders may reflect a shift toward increased functional specialization across hemispheres. This interpretation is supported by theories of hemispheric lateralisation in affective regulation, particularly the right hemisphere’s predominant role in processing negative emotions and autonomic arousal, and the left hemisphere’s association with approach-related affective states (20, 21). The administration of antidepressant medication has been demonstrated to induce a rebalancing of such activity, which may manifest as increased functional asymmetry. In healthy individuals, emotional regulation is characterised by efficient lateralised processing in the temporal lobes, regions critical for emotional memory, semantic integration, and social-emotional decoding. Consequently, a shift towards asymmetric connectivity may indicate a restoration of normative processing hierarchies disrupted in MDD. It is therefore hypothesised that responders may be regaining hemispheric specialisation necessary for adaptive affective responses (10–12).

#### D.2. Increase in symmetry among non-responders

The tendency toward increased symmetry (or lack of a decrease) observed among non-responders may reflect reduced inter-hemispheric differentiation, a phenomenon linked to inefficient or inflexible neural communication. One hypothesis is bilateral over, engagement of homologous temporal regions, potentially a compensatory response to impaired lateralised processing. This may reflect a rigid or maladaptive network configuration unable to support the neuroadaptive processes necessary for symptom improvement. Non-responders to antidepressants have been shown to exhibit reduced network flexibility and lower connectivity entropy (48–50), suggesting a form of cortical “inertia” during treatment. Resting-state fMRI in treatment-resistant depression likewise reports aberrant bilateral synchronisation, particularly in midline and temporal regions, arguably pathological redundancy rather than efficient lateralisation (51). EEG studies echo this with findings of increased bi-hemispheric coherence in neuropsychiatric states with poor prognosis, suggesting non-selective cortical recruitment (52).

#### D.3. Temporal-lobe specificity

The temporal lobes play crucial roles in semantic memory, affective processing, auditory language functions, and autobiographical recall, all processes commonly disrupted in depression. The right temporal cortex is especially implicated in emotional prosody and social context, whereas the left temporal lobe is more involved in language and semantic memory (53). fMRI shows lateralised functional disruptions in the DMN and temporal cortex in MDD, particularly among non-responders (54, 55). Recent EEG evidence also suggests that depressed patients exhibit reduced functional segregation, especially in the beta range (56).

The observed divergence in temporal symmetry between responders and non-responders likely reflects differential trajectories of functional reorganisation. Decreased symmetry in responders may signal a re-establishment of hemispheric specialisation required for adaptive emotional processing, whereas increased symmetry among non-responders could denote reduced network flexibility or compensatory bilateral engagement.

Taken together, these effects might highlight a temporally lateralized modulation of cross-hemispheric connectivity accompanying clinical improvement. Importantly, our symmetry findings do not contradict prior frontal or cingulate accounts; rather, they point to distributed, cross-hemispheric temporal synchronization changes that accompany early clinical improvement and may integrate with rACC/sgACC-centric predictors into a multi-marker panel (17). Moreover, while frontal asymmetry has faced replication challenges in recent meta-analyses (13, 14), our approach offers a computationally straightforward alternative that captures different aspects of network dysfunction. Practically, such network-level indices are inexpensive to compute from routine EEG and, as shown here, yield stable, directionally consistent group-level effects. Future work should explore whether combining temporal symmetry with frontal or cingulate markers improves prediction accuracy. Our analyses focused on temporal lobe defined regions and demonstrated convergent effects across both a core and an extended temporal mask. We did not aim here to determine whether similar asymmetry patterns might also be detectable at more global hemispheric scales, and systematic comparisons with whole-hemisphere or non-temporal masks are therefore an important direction for future work.

### E. Granularity of treatment response classification

An important consideration is that the binary classification of patients as responders or non-responders, based solely on MADRS and CGI scores, might oversimplify the complex nature of treatment responses. The dichotomous approach does not fully account for the varying degrees of improvement observed in clinical practice. For instance, partial responders, patients who show moderate but insufficient improvement, are not distinctly categorized, which limits our understanding of the nuances in treatment efficacy.

Future research could benefit from adopting a more granular classification system that accounts for varying levels of response, such as categorizing patients into full responders, partial responders, and non-responders. Such a system would enable researchers to explore the factors driving these different levels of response. This gradation could also reveal subtle but important trends in the data, such as the impact of specific parameters or treatment durations on partial responders. Additionally, incorporating intermediate categories may lead to more targeted treatment strategies and facilitate early intervention for patients unlikely to achieve full response. From a clinical perspective, a detailed stratification of treatment responses may help clinicians tailor treatments more effectively and improve long-term patient well-being.

### F. Predictive utility of early EEG markers

Quantitatively, early EEG-based response prediction is feasible but remains challenging and highly dependent on cohort characteristics and validation design. A machine-learning meta-analysis of EEG prediction studies in MDD reported pooled performance of 83.93% accuracy (95% CI: 78.90–89.29) and AUC = 0.850 (95% CI: 0.747–0.890) across studies (23); however, pooled estimates aggregate heterogeneous pipelines and should be interpreted as a benchmark rather than an expected value for any single dataset or feature family. Time-series-driven representations can also improve separability in some settings: for example, a motif-discovery framework applied to day-7 EEG reported accuracy of 0.731 (F1 = 0.722) in the *α* band for response prediction (24). In the present cohort, temporal symmetry change scores derived from longitudinal connectivity (Visit 2 − Visit 1) yielded modest but reproducible out-of-sample separability under repeated nested cross-validation (best-band median AUC ≈ 0.69), indicating above-chance discrimination but not a stand-alone clinical decision tool. We therefore view these symmetry measures as an interpretable, low-dimensional feature family that may complement established EEG predictors and richer time-series representations in future multi-marker models. External validation in independent multi-site cohorts will be essential before translational use.

It is also important to contextualize the observed performance within the broader EEG-based response-prediction literature. Reported accuracies and AUCs vary widely across studies as a function of sample size, outcome definition, and (critically) validation design, with externally validated performance typically lower than within-cohort estimates. Meta-analytic work on EEG/QEEG biomarkers underscores both the promise of electrophysiological predictors and the prevalence of heterogeneity and validation gaps (13). In this setting, the present results should be viewed as an interpretable, connectivity-level complement to established EEG markers rather than as a replacement for them.

However, the potential role of EEG changes beyond the first week warrants further investigation. While our study primarily emphasized early EEG dynamics, later changes may also contribute to understanding long-term treatment trajectories. This raises questions about whether these early improvements reflect intrinsic properties of the treatment timeline or whether longer durations of treatment elicit distinct neurophysiological patterns. Exploring these aspects could lead to a more nuanced understanding of how EEG signals evolve over time, shedding light on the interplay between treatment duration, neuroplasticity, and clinical outcomes. By examining these variables in future studies, we can uncover whether extending EEG monitoring throughout the treatment course offers additional predictive power and whether tailored interventions can optimize recovery for individual patients.

### G. Influence of treatment duration and demographic factors

The influence of treatment duration remains a largely unexplored yet critical factor. Investigating the interplay between age, treatment length, and outcomes could significantly refine clinical decision-making. For instance, understanding whether younger patients respond more quickly to shorter treatment regimens or whether older individuals require prolonged interventions could enable more personalized approaches to care. Future studies aimed at systematically evaluating these variables across diverse patient cohorts are essential to uncover the nuances of demographic and treatment-related influences on recovery trajectories.

Beyond their statistical contributions, incorporating demographic features like age and sex highlights the importance of designing models that are both inclusive and adaptive. By reducing potential biases, these features contribute to the robustness of predictive frameworks, laying the groundwork for further exploration in applied clinical settings. However, future research should expand on this foundation by exploring how other demographic or clinical variables, such as comorbid conditions or baseline symptom severity, may interact with EEG-derived features. Such investigations could deepen our understanding of individual variability in treatment response, enabling more precise stratification of patients and enhancing the overall efficacy of predictive models in psychiatry. Although responders and non-responders did not differ significantly in age, sex, or baseline MADRS scores (Table 3), small between-group differences in these clinical characteristics remain and cannot be entirely excluded as a source of residual confounding. In addition, the present cohort included heterogeneous treatment regimens and a range of illness durations, which we did not analyse in detail at the subgroup level; these factors may contribute to residual variability in both clinical outcome and EEG measures and should be examined more systematically in future work.

## Conclusion

Major Depressive Disorder (MDD) remains a pressing global health challenge, with substantial variability in how individuals respond to treatment. This variability underscores the urgent need for early biomarkers that can guide treatment decisions and reduce the time spent on ineffective therapies. In this study, we focused on symmetry features derived from EEG recordings of the temporal lobe, examining whether these features capture treatment-related changes in functional brain connectivity.

Using a graph-theoretical framework, we introduced both region-wise and electrode-wise symmetry measures based on lagged coherence, computed across standard EEG frequency bands. Our statistical analysis revealed consistent group-level differences between responders and non-responders in the *α, β*_1_, and *β*_2_ bands. In particular, we found that symmetry tended to decrease in the responder group and increase in the non-responder group following treatment. This directional pattern was observed across both binary and weighted versions of the features, suggesting a robust trend in temporal lobe dynamics.

To assess the potential utility of these features in predictive contexts, we applied them within a standard classification pipeline. While the resulting performance was moderate, it was stable across cross-validation folds, and certain symmetry features were repeatedly selected. This supports their relevance as discriminative signals, even if further development and validation are needed for clinical applications.

Our work contributes to a growing literature on EEG biomarkers by focusing on interhemispheric symmetry, an often underexplored aspect of brain connectivity despite its known relevance to emotional regulation and depression. The observed group-level differences suggest that monitoring changes in temporal lobe symmetry may help distinguish treatment responders from non-responders. By emphasizing interpretability and computational simplicity, the proposed features offer a useful foundation for future research in precision psychiatry.

Future work could expand this framework to include source-level EEG estimates or multimodal data integration. The proposed features offer a transparent and low-complexity approach that may support future developments in data-driven mental health research.

## Acknowledgments

This research was supported by the Czech Science Foundation [grant number GF21-14727K] and by the Czech Academy of Sciences, Praemium Academiae awarded to M. Paluš. The research of M. Brunovský was supported by the European Regional Development Fund Project Brain dynamics (Project No. CZ.02.01.01/00/22_008/0004643) and Charles University Cooperatio Neurosciences research program.

## Data availability

The data that support the findings of this study may be available upon reasonable request.

TC:ignore the command above ignores this section for word count

## Supplementary Note 1: Appendices

**Table 10:** Baseline (Visit 1) Mann-Whitney U tests comparing responders and non-responders (responders *n* = 84, non-responders *n* = 92) for all symmetry features across frequency bands. Entries are BY-adjusted *p* (*q*_BY_). None reached significance after correction.

| Feature | $\delta$ | $\theta$ | $\alpha$ | $\beta_1$ | $\beta_2$ | $\gamma$ |
| --- | --- | --- | --- | --- | --- | --- |
| RSTcore_bin | 1.000 | 1.000 | 1.000 | 1.000 | 1.000 | 1.000 |
| RSTcore_wei | 1.000 | 1.000 | 1.000 | 1.000 | 1.000 | 1.000 |
| RSText_bin | 1.000 | 1.000 | 1.000 | 1.000 | 1.000 | 1.000 |
| RSText_wei | 1.000 | 1.000 | 1.000 | 1.000 | 1.000 | 1.000 |
| ES_T3/4_bin | 1.000 | 1.000 | 1.000 | 1.000 | 1.000 | 1.000 |
| ES_T3/4_wei | 1.000 | 1.000 | 1.000 | 1.000 | 1.000 | 1.000 |
| ES_T5/6_bin | 1.000 | 1.000 | 1.000 | 1.000 | 1.000 | 1.000 |
| ES_T5/6_wei | 1.000 | 1.000 | 1.000 | 1.000 | 1.000 | 1.000 |

**Table 11:** Significant results of between-group Mann-Whitney U tests on Δ = V2 − V1 across features and frequency bands. Columns show BY-adjusted *q*_BY_, medians of Δ for responders and non-responders, and their difference (Δ_*N*_ − Δ_*R*_).

| Feature | Band | $q_{BY}$ | Median $\Delta_R$ | Median $\Delta_N$ | $\Delta_N - \Delta_R$ | Direction |
| --- | --- | --- | --- | --- | --- | --- |
| RSTcore_bin | $\alpha$ | 0.0015 | -0.1353 | +0.1392 | +0.2745 | ↓ Responders |
| RSTcore_bin | $\beta_1$ | 0.0108 | -0.0953 | +0.0497 | +0.1449 | ↓ Responders |
| RSTcore_bin | $\beta_2$ | 0.0276 | -0.0675 | +0.0156 | +0.0831 | ↓ Responders |
| RSTcore_wei | $\alpha$ | 0.0045 | -0.1677 | +0.2299 | +0.3975 | ↓ Responders |
| RSTcore_wei | $\beta_1$ | 0.0466 | -0.1100 | +0.0204 | +0.1304 | ↓ Responders |
| RSText_bin | $\alpha$ | 0.0119 | -0.2198 | +0.0958 | +0.3156 | ↓ Responders |
| RSText_bin | $\beta_1$ | 0.0021 | -0.1463 | +0.1574 | +0.3037 | ↓ Responders |
| RSText_wei | $\alpha$ | 0.0161 | -0.2880 | +0.2359 | +0.5239 | ↓ Responders |
| RSText_wei | $\beta_1$ | 0.0103 | -0.2185 | +0.1469 | +0.3654 | ↓ Responders |
| RSText_wei | $\beta_2$ | 0.0161 | -0.1072 | +0.0698 | +0.1770 | ↓ Responders |
| ES_T3/4_bin | $\beta_1$ | 0.0096 | -0.1675 | +0.2749 | +0.4424 | ↓ Responders |
| ES_T3/4_wei | $\beta_1$ | 0.0465 | -0.2972 | +0.1047 | +0.4020 | ↓ Responders |
| ES_T5/6_bin | $\alpha$ | 0.0147 | -0.4332 | +0.3428 | +0.7760 | ↓ Responders |
| ES_T5/6_bin | $\beta_1$ | 0.0018 | -0.4804 | +0.2576 | +0.7380 | ↓ Responders |
| ES_T5/6_wei | $\alpha$ | 0.0079 | -0.6720 | +1.1499 | +1.8220 | ↓ Responders |
| ES_T5/6_wei | $\beta_1$ | 0.0079 | -0.6846 | +0.2992 | +0.9838 | ↓ Responders |

**Table 12:** Post-treatment (Visit 2) Mann-Whitney U tests comparing responders and non-responders across features and frequency bands. Entries are BY-adjusted *p* (*q*_BY_).

| Feature | $\delta$ | $\theta$ | $\alpha$ | $\beta_1$ | $\beta_2$ | $\gamma$ |
| --- | --- | --- | --- | --- | --- | --- |
| RSTcore_bin | 1.000 | 1.000 | 0.011 | 0.017 | 0.880 | 0.927 |
| RSTcore_wei | 1.000 | 1.000 | 0.011 | 0.041 | 0.396 | 1.000 |
| RSText_bin | 1.000 | 1.000 | 0.039 | 0.006 | 0.951 | 1.000 |
| RSText_wei | 1.000 | 1.000 | 0.043 | 0.008 | 0.705 | 1.000 |
| ES_T3/4_bin | 1.000 | 1.000 | 0.367 | 0.008 | 1.000 | 1.000 |
| ES_T3/4_wei | 1.000 | 1.000 | 0.509 | 0.031 | 1.000 | 1.000 |
| ES_T5/6_bin | 1.000 | 1.000 | 0.157 | 0.089 | 1.000 | 1.000 |
| ES_T5/6_wei | 1.000 | 1.000 | 0.187 | 0.080 | 0.837 | 1.000 |

**Table 13:** Linear mixed-effects model estimates for group × time interactions across features and frequency bands. Each coefficient (*β*_interaction_) represents the between-group difference in change over time (difference-in-differences estimand); BY-adjusted *p*-values are reported as *q*_BY_.

| Feature | Band | $\beta_{\text{interaction}}$ | $q_{\text{BY}}$ |
| --- | --- | --- | --- |
| RSTcore_bin | $\delta$ | 0.001561 | 1.0000 |
| RSTcore_bin | $\theta$ | -0.091407 | 0.4279 |
| RSTcore_bin | $\alpha$ | -0.263508 | 0.0027 |
| RSTcore_bin | $\beta_1$ | -0.258364 | 0.0019 |
| RSTcore_bin | $\beta_2$ | -0.147444 | 0.0110 |
| RSTcore_bin | $\gamma$ | -0.049918 | 1.0000 |
| RSTcore_wei | $\delta$ | -0.000201 | 1.0000 |
| RSTcore_wei | $\theta$ | -0.100686 | 0.4606 |
| RSTcore_wei | $\alpha$ | -0.620674 | 0.0257 |
| RSTcore_wei | $\beta_1$ | -0.420475 | 0.0297 |
| RSTcore_wei | $\beta_2$ | -0.449117 | 0.0191 |
| RSTcore_wei | $\gamma$ | -0.008814 | 1.0000 |
| RSText_bin | $\delta$ | 0.014472 | 1.0000 |
| RSText_bin | $\theta$ | -0.116162 | 0.7300 |
| RSText_bin | $\alpha$ | -0.360953 | 0.0190 |
| RSText_bin | $\beta_1$ | -0.415231 | 0.0013 |
| RSText_bin | $\beta_2$ | -0.221439 | 0.0325 |
| RSText_bin | $\gamma$ | -0.048371 | 1.0000 |
| RSText_wei | $\delta$ | 0.017598 | 1.0000 |
| RSText_wei | $\theta$ | -0.107528 | 0.9043 |
| RSText_wei | $\alpha$ | -0.963905 | 0.0324 |
| RSText_wei | $\beta_1$ | -0.666336 | 0.0304 |
| RSText_wei | $\beta_2$ | -0.708423 | 0.0220 |
| RSText_wei | $\gamma$ | -0.068677 | 1.0000 |
| ES_T3/4_bin | $\delta$ | 0.003775 | 1.0000 |
| ES_T3/4_bin | $\theta$ | -0.369281 | 0.3335 |
| ES_T3/4_bin | $\alpha$ | -0.618101 | 0.2331 |
| ES_T3/4_bin | $\beta_1$ | -0.881060 | 0.0045 |
| ES_T3/4_bin | $\beta_2$ | -0.451566 | 0.0773 |
| ES_T3/4_bin | $\gamma$ | -0.149052 | 1.0000 |
| ES_T3/4_wei | $\delta$ | -0.001752 | 1.0000 |
| ES_T3/4_wei | $\theta$ | -0.290084 | 0.6090 |
| ES_T3/4_wei | $\alpha$ | -1.965858 | 0.1479 |
| ES_T3/4_wei | $\beta_1$ | -1.124379 | 0.0923 |
| ES_T3/4_wei | $\beta_2$ | -1.052408 | 0.0727 |
| ES_T3/4_wei | $\gamma$ | -0.618171 | 1.0000 |
| ES_T5/6_bin | $\delta$ | -0.023911 | 1.0000 |
| ES_T5/6_bin | $\theta$ | -0.290743 | 0.4734 |
| ES_T5/6_bin | $\alpha$ | -0.801515 | 0.0271 |
| ES_T5/6_bin | $\beta_1$ | -0.997680 | 0.0022 |
| ES_T5/6_bin | $\beta_2$ | -0.498634 | 0.0271 |
| ES_T5/6_bin | $\gamma$ | 0.072917 | 1.0000 |
| ES_T5/6_wei | $\delta$ | 0.009104 | 1.0000 |
| ES_T5/6_wei | $\theta$ | -0.323290 | 0.4207 |
| ES_T5/6_wei | $\alpha$ | -2.926048 | 0.0210 |
| ES_T5/6_wei | $\beta_1$ | -1.675937 | 0.0247 |
| ES_T5/6_wei | $\beta_2$ | -1.340052 | 0.0210 |
| ES_T5/6_wei | $\gamma$ | 0.373328 | 1.0000 |

